# Interplay between fecundity, sexual and growth selection on the spring phenology of European beech (*Fagus sylvatica* L.)

**DOI:** 10.1101/2023.04.27.538521

**Authors:** Sylvie Oddou-Muratorio, Aurore Bontemps, Julie Gauzere, Etienne Klein

## Abstract

**Background:** Plant phenological traits such as the timing of budburst or flowering can evolve on ecological timescales through response to fecundity and viability selection. However, interference with sexual selection may arise from assortative mating. This study aims to investigate how these three components of selection on spring phenology may combine in European beech populations in contrasting environments (high versus low altitude).

**Methods:** we monitored the timing of budburst (TBB) in 339 adult beech trees and estimated their fecundity using spatially explicit mating models. Fecundity selection was infered by regressing fecundities on TBB, while sexual selection was inferred by regressing fecundities on mating opportunities (i.e., TBB mismatch). The correlation between mates for flowering time (i.e., assortative mating) was estimated based on paternity analyses. Morever, TBB and growth were surveyed in 3261 seedlings from 40 families grown planted in a common garden, and viability selection was inferred by regressing growth on TBB.

**Results:** Overall, directional fecundity selection on female fitness favored trees with earlier TBB. Sexual selection acted only on male fitness through assortative mating favoring trees with mean TBB value (stabilizing selection). In the common garden, early budburst was associated with higher seedling growth. The respective intensities of directional and stabilizing selection varied with the environment: at low altitude, directional selection for earlier phenology was modulated by strong assortative mating and by an interaction effect between TBB an size on female fecundity, whereas at high altitude, directional selection for earlier phenology was reinforced by selection through male fecundity.

**Discussion:** This study showed that selection through female fecundity and seedlings growth predominantly selected for earlier TBB, while sexual selection on male fitness through assortative mating modulated this trend. This interplay between fecundity and sexual selection calls for an integrative approach to predict the evolution of spring phenology under a changing climate.

## Introduction

During the last few decades, many changes in phenology (i.e., the timing of biological events) were observed and attributed to climate change (Parmesan and Yohe 2003). In particular, records of leafing, flowering and fruiting have advanced significantly in temperate zones (Menzel et al. 2006), consistent with the increase of spring/summer temperatures. Rapid evolution of phenological traits in response to selection has been reported (Franks et al. 2007; Hamann et al. 2018). However, it is still largely unknown to what extent evolution over a few generations may contribute to the response of plant populations to ongoing climate change (Merilä and Hendry 2014). Moreover, in many plants, vegetative phenology (the timing of germination, stem and leaf development) and reproductive phenology (the timing of flowering and fruiting) are tightly synchronized throughout the yearly cycle. Hence, selection on phenological traits is likely to result from a complex interplay between viability selection (selection for phenotypes that increase survival), fecundity selection (selection for phenotypes that increase fecundity) and sexual selection (selection arising from competition for mating partners or their gametes) (Figure 1). This study aims to investigate selection on spring phenology in a temperate tree species along an altitudinal gradient while into account these different components of selection.

**Figure 1.**
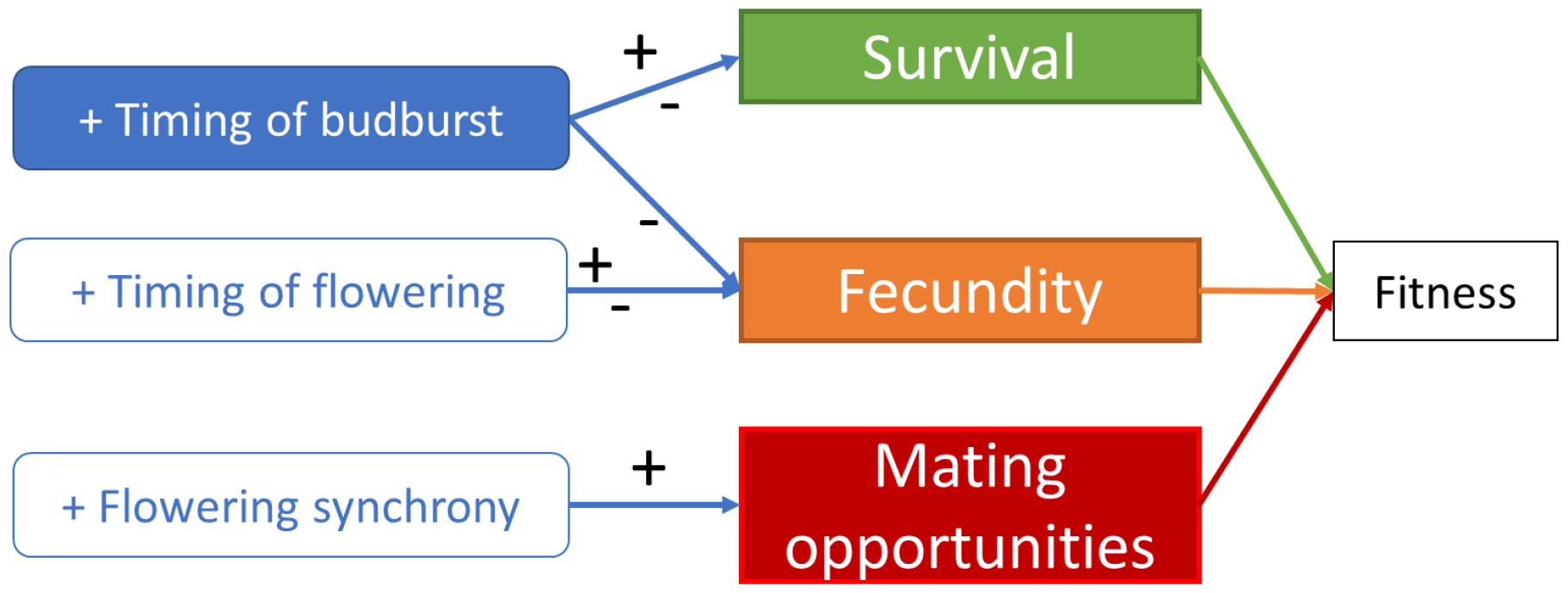
A schematic representation of the expected relationships between vegetative/flowering phenology and fitness at the individual level. Within a population, individuals with delayed timing of budburst relative to the population mean are expected to have higher survival through frost avoidance, but also lower fecundity and survival through reduced length of the growing season, and hence reduced reserves. Individuals with delayed timing of flowering relative to the population mean are expected to have higher fecundity because more resources can be accumulated and invested in reproduction, however at the cost of reduced time for seed maturation. Finally, synchronized flowering with the other individuals in the population (i.e., assortative mating) is expected to maximize the number of mates. The combination of these different components of selection determines the optimal values of TBB and flowering times, i.e., those maximizing fitness. Note that selection and hence the optimal values of phenological traits can also vary between environments. Colored boxes represent traits for which we have observations/estimations in this study. The sign “+” (respectively “-”) indicates an increase (respectively decrease) in the value of the variable under consideration.

Most selection studies on the timing of flowering have been conducted in short-lived herbaceous plants (Geber & Griffen, 2003; Munguía-Rosas et al., 2011), while the adaptive value of vegetative phenology has been mainly investigated in long-lived forest trees (Alberto et al. 2013). In both cases, stabilizing selection is the most straightforward expectation within-population, considering life history in temperate ecosystems. In the former case, this is because the fecundity benefits of flowering early (sufficient time for seed maturation) are expected to balance with those of flowering later, as early reproduction usually entails reproducing at a small size with limited resources available for offspring production. Note that this expectation could be different for long-living plants for which resources have been accumulated previous year (Hacket-Pain et al. 2018). Yet, early flowering plants are generally found to be favored (Geber and Griffen 2003; Munguía-Rosas et al. 2011), an apparent paradox for which different explanations have been proposed (Austen et al. 2017).

Regarding vegetative phenology in long lived plants inhabiting temperate ecosystems, stabilizing selection is expected to be driven by the balance between the benefits of: (1) emerging leaves later and avoiding frost damages on vegetative and reproductive organs, especially in early spring (Augspurger 2013; Bigler and Burgmann 2018); and (2) emerging leaves earlier and maximizing the duration of the growing season, which determines the resource level acquired by photosynthesis (Keenan et al. 2014; Richardson et al. 2006). More complex situations may occur when other abiotic or biotic stresses are considered (e.g. early flushing may amplify drought effects, Meier et al. 2021). Common-garden experiments generally demonstrate significant genetic differentiation of phenological traits between tree provenances along environmental gradients, suggesting that the differences in climatic conditions led to the evolution of different phenological schedules contributing to populations’ local adaptation (Alberto et al. 2013). However, experimental selection studies on tree vegetative phenology remain limited in comparison with those on plant flowering phenology (but see Bontemps et al. 2017; Alexandre et al. 2020; Westergreen et al. 2023). A recent simulation study with a process-based phenological model accounting both for fecundity and viability selection predicted selection towards earlier TBB across a climatic gradient, and realized TBBs always later than the value conferring highest fitness in different tree species (Gauzere et al., 2020). Moreover, these simulations showed that the strength of this selection was stronger at high than low altitude, i.e., in the conditions where the growing season is more limiting for the maturation of fruits.

Compared to fecundity or viability selection, the role of sexual selection on the evolution of phenology remains understudied, even though the existence of sexual selection in plants is now widely acknowledged (Moore and Pannell 2011). Yet, assortative mating for flowering phenology, that is the positive correlation between male and female flowering time across mated pairs, is obligate in plants (Weis et al. 2014). Hence, variation of individual flowering phenologies within the population may result in sexual selection. Moreover, phenological assortative mating is by nature density-dependent, as any individual synchronized with the rest of the population will gain opportunities for mating (Weis et al. 2005). Hence, assortative mating is expected to generate a form of stabilizing sexual selection towards an optimal timing of flowering maximizing mating opportunities (Soularue et al. 2023). Finally, due to anisogany (the higher cost of producing female versus male gametes), male reproductive success is generally expected to be more limited by mating opportunities than than by investment in each gamete, whereas female reproductive success should depend on their ability to produce viable ovules and seeds rather than on the probability of having ovules fertilized (one of Bateman’s principles; Bateman 1948; Tonnabel, David, & Pannell, 2019). These contrasting challenges could lead to different patterns of selection on phenology through male and female reproductive functions.

Distinguishing fecundity from sexual selection on phenological traits may be particularly challenging, as both jointly act within a single reproduction episode. However, while fecundity selection can occur even under unlimited access to mates, sexual selection involves limited mating opportunities. Hence, the relationship between phenology and fitness (e.g., phenotypic selection analyses, Lande and Arnold 1983) is considered to inform about the joined effects of fecundity and sexual selection (i.e., natural selection), while the relationship between phenology-related mating opportunities and fitness (e.g., Bateman’s gradient analyses, Bateman 1948) informs about sexual selection on phenology.

This study takes advantage of the extensive physiological knowledge on a major monoecious tree, the European beech, and of a well-studied altitudinal gradient in South-Eastern France, to estimate different types of selection on the timing of budburst (TBB). European beech is an early flushing deciduous species (Davi et al. 2011), sensitive to frost damages (Lenz et al., 2013). Along the studied gradient, vegetative phenology was monitored both *in situ* and *ex situ*, in a common garden of maternal progenies (Oddou-Muratorio et al. 2021). Previous studies showed that, *in situ*, budburst occurs ~9.8 days earlier at the lower altitude plot compared to the upper altitude plot (Davi et al. 2011), but that, in the common garden, the lower plot is ~2.1 days late compared to the upper plot (Gauzere et al., 2020). This is a classical counter-gradient pattern where the *in situ* plastic response of TBB to different temperature accumulation at the two altitudes (Table 1) hides the genetic differentiation revealed in the common garden (Gauzere et al., 2020). Phenotypic selection analyses conducted at the lower plot found that growth and reproductive (seed set) performances could be maximized either by a water-uptake strategy, including early budburst, or by a water-saving strategy, including late budburst (Bontemps et al. 2017). Finally, male and female fecundities were estimated for all the adults in the lower and upper plots through paternity or parentage analysis of germinated seeds and established saplings (Oddou-Muratorio et al., 2018), which showed that both female and male fecundities increased with tree size and decreased with density and competition in the neighbourhood, the details of these effect varying among plots at different altitude.

**Table 1:** Climatic context and main expectations regarding selection on phenology at the two studied plots. Climate is synthetised by six variables computed from the long-term daily dataset from 1959 to 2013 described in Davi & Cailleret (2017): the mean annual temperature (tmean, °C), the maximum temperature of July (tmax, °C), the minimum temperature of January (tmin, °C), the sum of growing degree days (GDD, °C), the number of frost days, the water stress level between May and September (mm/m2/day), computed as the difference between ETP and precipitations. See Fig S1 for additionnal details.

| Plot | Climate |  |  |  |  |  | Main constraint | Expectation |
| --- | --- | --- | --- | --- | --- | --- | --- | --- |
|  | tmean | tmax | tmin | GDD | NFD | Stress |  |  |
| **N1-LOW** | 9 | 22.3 | -0.5 | 3060.9 | 14.3 | 154.7 | High water stress | Both early or late budburst may enhance survival and fecundity (Bontemps et al. 2017) |
| **N4-HIGH** | 6.3 | 18.5 | -2.8 | 2187.1 | 35.1 | 69.2 | Short growing season | Intense fecundity selection for early phenology is expected |

The specific aim of this study was to simultaneously investigate fecundity, sexual and viability selection on spring phenology. Our main hypothesis is that inter-individual variations in TBB are strongly correlated with inter-individual variations in the timing of flowering, making TBB an appropriate trait to study these different components of selection. First, we estimated fecundity selection by regressing male and female effective fecundity on TBB (both measured *in situ*). Second, we used paternity analyses to investigate assortative mating, and we estimated sexual selection by regressing male and female fecundities on mating opportunities, as measured by TBB mismatch within mating neighborhoods (also measured *in situ*). Third, we estimated viability selection in the community garden by analyzing the relationship between TBB and seedlings growth, under the hypothesis that vigor (i.e. growth capacity) is positively associated with viability (Collet and Le Moguedec 2007). For these three inferences of fecundity, sexual and viability selection, we relied on the classical metrics of selection gradients (the regression coefficients of relative fitness on a trait, Lande & Arnold, 1983). In addition, we analysed both the upper and lower plots along the altitudinal gradient, as these contrasting environments are expected to result in different selective constraints (Table 1).

## Methods

### Studied species and site, sampling design

The European beech is a monoecious, wind-dispersed, predominantly outcrossed tree species (Gauzere, Klein, & Oddou-Muratorio, 2013). Male and female flowers are borne on the same branches and open as the leaves unfold (Nielsen & Schaffalitzky de Muckadell, 1954; Packham et al., 2012), between April and May. Beech is protogynous, i.e. male flowers produce pollen after the peak of receptivity of the stigmas of the same plant (Nielsen and Schaffalitzky de Muckadell 1954).

Mont Ventoux is located at the warm and dry southern margin of the European beech distribution, and the climate is typical of low altitude mountains with Mediterranean influences (weather station of Mont Serein, 1 445 m a.s.l., 1993–2006; mean annual temperature of 6.8°C and mean annual rainfall of 1300 mm). On the northern face of Mont Ventoux, the beech forest ranges almost continuously from 750 to 1700 m above sea level. This steep altitude gradient provides almost linear variation in mean temperature and humidity with altitude (Cailleret and Davi 2011). We studied two plots at opposite positions along an altitudinal gradient, named N1-LOW (1.3 ha; 1,020 m a.s.l.), and N4-HIGH (0.8 ha; 1,340 m a.s.l.). N1-LOW is at the lower limit of the altitude range for European beech on Mont Ventoux, while N4-HIGH is at the upper limit for sexual reproduction.

In 2009, one large masting event occurred, which provided a unique opportunity to collect seeds and monitor regeneration. All potentially reproductive trees were mapped, measured and sampled for genetic analyses (164 at plot N1-LOW and 365 at plot N4-HIGH). Mother-trees were chosen among the trees with medium to high seed production, ensuring a minimal distance of 10 m between two mother-trees, and covering the whole plot area. Open-pollinated seeds were collected from 20 mother-trees at each plot (40 families for this study, among 60 in total), germinated and sown in a greenhouse. These open-pollinated seeds first allowed us to estimate patterns of pollen flow and male fecundity (see below). Moreover, the common-garden experiment was arranged in 50 complete blocks (with two seedlings per family per block, Gauzere et al., 2020) and divided in two contrasted experimental conditions: “watered” (from block 1 to 25) versus “water-stressed” (from block 26 to 50; Oddou-Muratorio et al. 2021). Briefly, these two conditions allow us to contrast situations of non-limiting *versus* limiting water availability, and to investigate the plastic response of traits to water stress, even though the levels of water stress experienced in the second condition do not match to those experienced in situ. Seedlings of the watered condition were analyzed in a quantitative genetic framework to investigate the within-family, among-families within-plot and among-plots components of the genetic variation at several functionnal traits (Gauzere et al., 2016, 2020). In this study, we took the opportunity to compare the open-pollinated seedlings from plots N1-LOW and N4-HIGH growing in the two experimental conditions.

Finally, in September 2010, we sampled in situ seedlings originating from the same reproduction event in 2009 and germinated in spring 2010 (223 seedlings at plot N1-LOW and 250 seedlings at plot N4-HIGH). These established seedlings allowed us to estimate patterns of seed flow and female fecundity (see below). Spring at year 2009 was not colder as an average year considering the mean and minimal temperatures from March to June (Fig. S1). Not late frosts (ie temperatures <-4°C after budburst) were observed in 2009 at any site.

### Phenology measurement in situ and ex situ (common garden)

In beech, the flowering phenology is hard to follow because (1) it occurs when leaves are spread out, (2) the succession of the flowering stages is rapid, and (3) the reproductive organs are small. However, as in oaks (Franjic et al., 2011), reproductive buds open very shortly after leafing (Nielsen and Schaffalitzky de Muckadell 1954). Therefore, we employed budburst phenology as a proxy of reproductive phenology.

The budburst was surveyed in situ in spring 2009 on 147 adult trees in population N1-LOW, and 192 adult trees in N4-HIGH. The budburst phenology was characterized using the five stages described by Davi *et al*. (2011) and Jean *et al*. (2023): 1) dormant buds; 2) swelling buds; 3) broken bud scales; 4) emerging leaves; 5) spread out leaves (Fig. S2). The phenological stages of each adult tree were noted on 15 different dates in population N1-LOW (between the 23^th^ of March and the 4^th^ of May 2009), and on 13 different dates in population N4-HIGH (between the 24^th^ of March and the 5^th^ of May 2009). At each date, individual stage of development was assessed globally for the upper and lower part of the crown, and then average into a single stage value. Then, a phenological score sum (PSS) was computed for each tree as the sum of the phenological stages observed over all of the dates: the higher the PSS at a given date of measurement, the earlier and quicker was leaf unfolding (Bontemps et al. 2017). We also used a linear interpolation to estimate the timing of budburst (TBB) as the date of passage (number of days since 1st January) from stage 2 to 3, stage 3 being one the most sensitive stage to frost damages. Finally, we computed the spread of budburst for each adult tree from the temporal sequence of phenological scores, as the number of days where the phenological stage was >2 and ≤4 (i.e., the duration of stage 3).

The budburst was surveyed ex situ for seedlings in the common garden using five stages (Gauzere et al., 2016). The phenological stages were noted on 4 different dates (between the 5th and 26th of April 2011). We used linear interpolation to estimate TBB as the date of passage (number of days since 1st January) from stage 2 to 3.

### Estimation of male and female fecundities

Male and female fecundities were estimated using spatially explicit mating models as described in Oddou-Muratorio et al. (2018). Briefly, thes models consider mating and dispersal events in a hermaphroditic plant population, and allows individual fecundities to be estimated together with mating system parameters, using genotypes and positions of potential parents and their offspring. It is implemented in a Bayesian framework in the MEMM software. First, the individual male fecundities were estimated with MEMM, jointly with the pollen dispersal kernel, the selfing rate and the pollen migration rate, from the open pollinated seeds with known mother tree. Second, female effective fecundities were estimated jointly with male fecundities, the pollen and seed dispersal kernels, the selfing rate and the pollen and seed migration rates, with another version of MEMM taking as input one-year established seedlings without any known parent.

Remarkably, MEMM estimates of fecundity account for the effect of the relative positions of putative parents and offspring, while at the same basic fecundity, putative parents closer to an offspring would have a higher parentage probability in uncorrected models. Hence, by using MEMM, estimates of fecundity are not sensitive to spatial biases due to sampling design, or edge effects. Moreover, MEMM estimates of fecundities are *effective*: male fecundity is a proxy of the effective amount of pollen achieving successful pollination, and female fecundity is a proxy of the effective number of seeds achieving successful germination and establishment in the population. Therefore, these estimates account for the individual effects (maternal or genetic) that modify the success of mating, including differences in pollen tube growth, seed abortion (for male fecundity) and in seed maturation, germination or early survival during the post-dispersal processes (for female fecundity). Effective fecundity provides more realistic estimates of individual plant contribution to the next generation than simpler estimates, such as fruit or seed set. Finally, MEMM estimates of fecundity are relative, and consider uncertainty in parentage reconstruction. Indeed, MEMM does not categorically assign parents to offspring, but rather consider the likelihood of all adults to be the parent of each offspring, accounting for the genotypes of adults and offspring and allowing genotyping errors.

Estimations were performed separately on each plot. The MCMC procedure to estimate individual fecundities and mating system parameters is described in details in Oddou-Muratorio et al. (2018). For this study, we used only the fecundities estimated for those adult individuals for which vegetative phenology was monitored, that is: 147 among the 164 adults at plot N1-LOW, and 192 among the 365 adults at plot N4-HIGH.

### Fecundity selection analyses on adult trees, *in situ*

To investigate fecundity selection, we used selection gradient analysis (Lande and Arnold 1983), with MEMM estimates of fecundity as the response variable, and TBB as the predictor. Because fecundity variations are shaped by many other factors besides phenology, we included size and competition effects and their interactions using a hierarchical procedure, and selected the most parsimonious model to estimate the effect of phenology.

For each sex (male and female) and each plot (N1-LOW and N4-HIGH), seven hierarchical models were compared. We first fitted a baseline model M1 including only the predictor of interest (TBB):

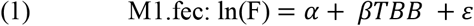

where F is the fecundity, α is the origin of the regression, β is the directional selection gradient on TBB, and ε is the residual.

In a previous study not including phenological traits (Oddou-Muratorio et al. 2018), we showed that both female and male fecundities increased with tree size and decreased with density and competition in the neighborhood. As selection probably simultaneously acts on these different correlated characters (phenology, size, competition), we fitted three models including size or/and competition variables, in addition to TBB:

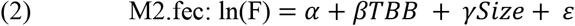

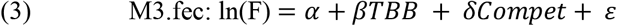

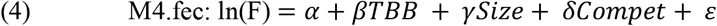

Note that several variables were used to measure size and competition (Table 2 and Oddou-Muratorio et al. 2018). Tree size was assessed by measuring the Diameter at Breast Height (Dbh), but as beech sometimes produces stump shoots resulting in multiple stems, we measured both the maximum Dbh (MaxDbh) and the sum of Dbh (SumDbh) of all the stems produced by a given genotype. Competition on each adult tree was assessed using (1) beech density in a radius of 20 m (ConDens20), (2) a competition index integrating the density and diameter of beech competitors in a radius of 20 m (ConMartin20) and (3) tree stature (a class variable with 3 levels: dominant, codominant, and suppressed). Based on the previous results of Oddou-Muratorio et al. (2018), we chose the most pertinent variable for each sex and plot (Table 2).

**Table 2:** Variables included in the fecundity selection analyses. The size and competition variables were identified as best predictors (BestPred) of female fecundity (*F♀*) and male fecundity (*F♂*) in the previous study of Oddou-Muratorio et al. (2018).

| Category | Variable name | Variable definition | BestPred | Range at N1-LOW | Range at N4-HIGH |
| --- | --- | --- | --- | --- | --- |
| Size | MaxDbh (cm) | Maximum diameter of the clonal copies | $F_{\text{♀}}$ | 10.1-45.4 | 9.2-28.3 |
| | SumDbh (cm) | Sum of diameters of the clonal copies | $F_{\text{♂}}$ | 11.6-231.6 | 10.2-127.1 |
| Competition | ConDens20 | Conspecific local density in a radius of 20 m | $F_{\text{♀}}$ | 2-50 | 6-115 |
| | TotMartin20 | Total competition index in a radius of 20 m | $F_{\text{♀}}$ and $F_{\text{♂}}$ | 8.7-100 | 43.0-127.4 |
| | Stature | dominant, codominant, or suppressed | $F_{\text{♂}}$ | 44, 51, 52, resp. | 35, 66, 91, resp. |

\* this variable was retained as the estimator of phenological mismatch in the fecundity and sexual selection analyses of this study.

Finally, we also fitted three models including two-way interaction terms between TBB and size/competition covariates, in order to account for possible changes in the relationship between TBB and fecundity depending on size or competition:

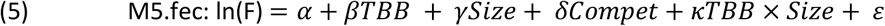

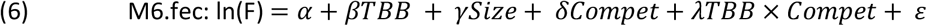

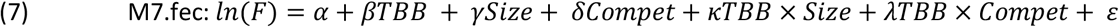

We compared these seven hierarchical models based on the Akaike information criterion (Akaike 1987) corrected for small sample size (AICc, Burnham and Anderson 2002), and we selected the most parsimonious model, denoted BestFec-Model in the following. Significance of the effects (AIC and *p*-values) was assessed with the function *drop1* of the R package *stats*.

A quadratic effect of TBB can also be included to estimate stabilizing selection through a bell-shaped response function (Lande and Arnold 1983). This was done by adding an additional term γTBB^2^ in the BestFec-Model selected above.

### Variable transformation and model fitting

Note that MEMM estimates of fecundity are relative, as required for selection gradient estimation (Lande and Arnold 1983). Moreover, the fecundities were log-transformed to approach Gaussian distribution and to account for the higher variance associated to higher fecundities. Besides, all the predictor variables (including TBB and PMis) were scaled to mean zero and unit variance. Such transformation of the predictor variables allows improving the interpretability and comparability of the estimated regression coefficients, especially when interactions are present (Schielzeth 2010). Once the best model selected, we estimated the standardized selection gradients on fecundity by fitting this selected best model without log-transformation of fecundity.

All models were fitted using the *lm* function implemented in R-base. Model comparisons was performed using the *aictab* function of the R package ‘AICcmodavg’ (Mazerolle 2020). For the best models, the residuals were visually inspected through a plot of residuals vs predicted. All the analyses are available as online supplementary material (file SelectionAnalyses_adult.html at https://doi.org/10.57745/ZVPNXX).

### Mating opportunities and assortative mating estimation

Direct observation of mating or pollination events being impossible in this anemophilous species, we used phenological mismatch as a proxy of mating opportunities. We computed the sum of phenological mismatches between each adult tree *i* and its neighbors in a radius R as: |*PMis*_*i*_|_*s*_ = ∑_*j in R*_|*TBB*_*i*_ − *TBB*_*j*_|, with R-values of 20, 50, 75 or 100 m. Note that similar |PMis|_s_ values can be obtained either with a low density and large asynchrony or with a high density and low asynchrony. We also computed a mean phenological mismatch |PMis|_m_, weighted by density. We hypothesize that the greater the phenological mismatch, the lower the opportunities of mating. Note also that using absolute mismatches implicitly assumes a symmetric effect asynchrony (earlier and later trees plays the same role for mating opportunities).

We estimated the strength of assortative mating as the correlation in vegetative phenologies between mates. We used the genetic data of maternal progenies (seedlings of the common garden) and adults trees to run paternity analyses and identify mating pairs, i.e. the most likely father siring a known mother. We used the genotypes of all the sampled adult trees in situ (147 trees at plot N1-LOW, and 192 trees at plot N4-HIGH) and of 1414 seedlings growing in the common garden (694 seedlings from plot N1-LOW and 720 seedlings from plot N4-HIGH, for an average ~35.3 seedlings per mother tree.

The genotypes of seeldings and adults were scored at a combination of 13 microsatellite loci (Oddou-Muratorio et al. 2018). The number of alleles observed in each cohort was greater than 106. Combine across all 13 loci, the exclusion probability of a non-father was > 0.9999 at both plots. Paternity assignments were conducted using the maximum-likelihood procedure implemented in the software CERVUS v.3.0.7 (Marshall et al. 1998; Kalinowski et al. 2007). Likelihood scores, based on allele frequencies in the experimental population, were calculated for each seed /potential father couple. To determine whether the paternity of each offspring could be assigned to the father with the highest likelihood, we used the difference in likelihood scores (ΔLOD) between the two most likely pollen donors. The critical value (ΔC) of ΔLOD below which paternity/parentage could not be assigned at 80% was determined using a distribution of Δ obtained from 5 000 simulated mating events. This distribution was generated using the following simulation parameters: 1% of genotyping error and no unsampled parents. Indeed, considering simultaneoulsy the risk of genotyping error and the subsampling of the breeding male population may inflate the lack of power in detecting the true father although it was sampled (type II error rate)(Oddou-Muratorio et al, 2003).

### Sexual selection analyses on adult trees, *in situ*

To investigate sexual selection, we used Bateman gradient analysis (Bateman 1948; Tonnabel et al. 2019), with a proxy of mating opportunities as predictor (here, phenological mismatch), and MEMM estimates of fecundity as the response variable.

We followed the same strategy and methods as described above for fecundity selection. For each sex (male and female) and plot (N1-LOW and N4-HIGH), we fitted seven models as described by equations (1) to (7), but replacing TBB by PMis, the phenological mismatch between each tree and its neighbors in a 20 m radius. For instance, for the first model:

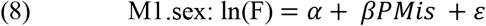

We compared the seven models based on the AICc, and selected the most parsimonious model, denoted BestSexSel-Model in the following.

We finally fitted a compound model, derived from the BestFec-Model but adding PMis as predictor, and we compared the BestFec-Model, the BestSex-Model, and the compound model.

### Seedlings growth measurements

We measured seedlings diameter (D2011_start_, D2011_end_) and height (H2011_start_, H2011_end_) in the common garden on two dates (respectively April 2011, and September 2011). This allowed us to estimate diameter growth in 2011 as GrowthD = D2011_end_ - D2011_start_ and height growth in 2011 as H2011_end_ - H2011_start_. In total, growth was measured in 2011 for 1552 seedlings originating from 20 families at plot N1-LOW, and for 1709 seedlings originating from 20 families at plot N4-HIGH. These seedlings were grown either in “watered” condition (from block 1 to 25, 1652 seedlings) or in “water-stressed” condition (from block 26 to 50, 1609 seedlings).

### Growth selection analyses on seedlings, in the common garden

As the common garden was designed to minimize seedlings mortality, we focused on growth as a performance trait related to viability, which is particularly expected when competition is homogeneous among seedlings (Collet and Le Moguedec 2007). We used the following mixed model to investigate the effect of phenology and plot on annual growth in diameter and height (respectively *GrowthD* and *GrowthH*) during year 2011:

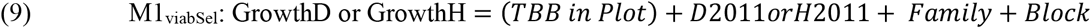

Where D2011 (respectively H2011) is the initial diameter (respectively height) in spring 2011, introduced to account for difference of vigor among seedlings. We tested for the effect of TBB nested within plot (N1-LOW or N4) to account for the fact that the effect of TBB on growth may differ among plots (knowing moreover that TBB is on average higher for seedlings at plot N1-LOW as compared to plot N4, Gauzere et al. 2020a). Family (the maternal family of the seedlings) and Block (the trial unit to which seedlings belongs to) were introduced as random factors to remove undesirable variation in growth related respectively to genetic variation for phenology and microenvironmental effects (e.g., half the bocks received a water-stress treatment).

We also tested another mixed model:

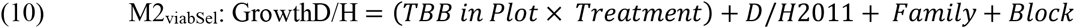

where the treatment (watered vs water-stress) was specified as a fixed effect, in order to investigate whether the effect of TBB on growth may differ among plots and among treatments.

These models were fitted with the function *lmer* in *lme4* package (Bates et al. 2015). All the analyses are available as online supplementary material (file GrowthSelectionAnalysis_seedlings.html at https://doi.org/10.57745/ZVPNXX).

## Results

### Preliminary examination of interindividual variations in phenology and fecundity

The timing of budburst (TBB) was observed to spread over 17 days at plot N1-LOW, with a mean TBB on April 20^th^ (Fig. S3). At plot N4-HIGH, TBB was observed to spread over 13 days, with a mean TBB on May 4th. Plot N4-HIGH showed a smaller inter-tree variance of TBB, with a significant proportion of trees having the same TBB of 124. Larger trees at plot N1-LOW had an earlier budburst (corr_TBB-circ_ = −0.15, p-value=0.007), while there was no significant relationship between size and TBB at plot N4-HIGH (corr_TBB-circ_ = −0.02, p-value=0.12). The spread of budburst within a tree was greater at plot N1-LOW (mean spread = 4.8 days) than at plot N4-HIGH (mean spread=2.9 days, see Fig. S4A). Trees with later budburst also exhibited a greater spread of budburst at plot N1-LOW (corr_TBB-spread_ = 0.57, p-value< 10^-3^), but not at plot N4-HIGH (corr_TBB-spread_ = 0.09, p-value=0.20; see Fig. S4B).

Male fecundities, as estimated by MEMM, followed a strongly L-shaped distribution (Fig. S5A). At plot N1-LOW, male fecundities (MF) ranged from 0.013 to 10.5 (median = 0.33) and 97 trees (66%) exhibited a non-negligible male fecundity. At plot N4-HIGH, male fecundities ranged from 2.10^-3^ to 16.6 (median = 0.016) and 69 trees (36%) exhibited a non-negligible male fecundity. Female fecundities, as estimated by MEMMseedlings, also followed a strongly L-shaped distribution (Fig. S5B). At plot N1-LOW, female fecundities ranged from 3.10^-3^ to 13.3 (median = 0.014), and 55 trees (37%) exhibited a non-negligible female fecundity. At plot N4-HIGH, female fecundities ranged from 3.10^-4^ to 25.8 (median = 0.005), and 30 trees (16%) exhibited a non-negligible female fecundity.

### Fecundity selection analyses for TBB, based on adult trees in situ

The study demonstrated that earlier budburst had a positive effect on female fecundity. This was observed in all trees at high altitude and in larger trees at low altitude (Table 3). The best model for female fecundity at plot N1-LOW included TBB, size, competition and their interactions. Delayed TBB had a negative impact on female fecundity in the larger trees, but a positive impact on smaller trees, as illustrated by figure 2A. Delayed TBB had a significant negative effect on female fecundity for the more competed trees (Fig. 2B). The directional selection gradient estimated for TBB was marginally significant (β_TBB_ = −0.37, p = 0.07; Table 3), indicating a positive effect of earlier TBB on female fecundity for a tree with average DBH and average competition.

**Table 3.** Fecundity selection on female (*F*♀) and male (*F*♂) fecundities at plots N1-LOW and N4-HIGH. Fecundity selection on phenology was assessed through the effect of TBB on fecundity, accounting for joint effect of size and competition (see variable names in Table 1). We selected the most parsimonious regression model for each sex and plot (Table S2) to estimate the effect’s coefficient and Sum Of Squares (SOSq) associated with each term. *F*♀ and *F*♂ were log-transformed.

| Term | Coefficient | SofSq | AIC | p-value |
| --- | --- | --- | --- | --- |
| $F_{\text{♀}}$, plot N1-LOW | Adjusted $R^2 = 0.1414$; p-value $< 0.001$; $AIC_{\text{ref}} = 262.85$ | | | |
| TBB | -0.37 | 18.91 | 264.24 | 0.072 |
| MaxDbh | 0.78 | 61.06 | 271.53 | 0.001 |
| ConMartin20 | 0.06 | 0.36 | 260.91 | 0.802 |
| TBB:MaxDbh | -0.88 | 54.02 | 270.34 | 0.003 |
| TBB:ConMartin20 | -0.55 | 32.28 | 266.59 | 0.019 |
| $F_{\text{♀}}$, plot N4-HIGH | Adjusted $R^2 = 0.145$; p-value $< 0.001$; $AIC_{\text{ref}} = 370.43$ | | | |
| TBB | -0.34 | 21.97 | 371.09 | 0.071 |
| MaxDbh | 0.78 | 87.06 | 380.67 | 0.000 |
| ConDens20 | -0.37 | 19.26 | 370.68 | 0.091 |
| $F_{\text{♂}}$, plot N1-LOW | Adjusted $R^2 = 0.119$; p-value $< 0.001$; $AIC_{\text{ref}} = 112.01$ | | | |
| TBB | -0.08 | 0.88 | 110.44 | 0.519 |
| SumDbh | 0.56 | 45.31 | 130.54 | $< 0.001$ |
| $F_{\text{♂}}$, plot N4-HIGH | Adjusted $R^2 = 0.176$; p-value $< 0.001$; $AIC_{\text{ref}} = 336.80$ | | | |
| TBB | -0.41 | 31.78 | 340.50 | 0.019 |
| SumDbh | 0.46 | 33.64 | 340.83 | 0.015 |
| Stature | Dom : 0.31 | 96.33 | 349.60 | $< 0.001$ |
|  | Cod : 0.74 |  |  |  |
|  | Suppr : -1.05 |  |  |  |

**Table 4:** Sexual selection on female (*F*♀) and male (*F*♂) fecundities at plots N1-LOW and N4-HIGH. Sexual selection on phenology was assessed through the effect of phenological mismatch (PMis) on fecundity accounting for joint effect of size and competition. We selected the most parsimonious regression model for each sex and plot (Table S3) to estimate the effect’s coefficient and Sum Of Squares (SOSq) associated with each term. *F*♀ and *F*♂ were log-transformed.

| Term | Effect | SofSq | AIC | p-val |
| --- | --- | --- | --- | --- |
| $F_{\text{♀}}$, plot N1-LOW | Adjusted $R^2=0.088$; p-value<0.001; AIC<sub>ref</sub>=268.86 | | | |
| PMis | 0.13 | 2.34 | 267.25 | 0.536 |
| MaxDbh | 0.85 | 95.81 | 282.07 | <0.001 |
| $F_{\text{♀}}$, plot N4-HIGH | Adjusted $R^2=0.117$; p-value<0.001; AIC<sub>ref</sub>=373.91 | | | |
| PMis | -0.2908 | 16.119 | 374.27 | 0.128 |
| MaxDbh | 0.9382 | 167.782 | 395.14 | <0.001 |
| $F_{\text{♂}}$, plot N1-LOW | Adjusted $R^2=0.199$; p-value<0.001; AIC<sub>ref</sub>=98.04 | | | |
| PMis | -0.4436 | 28.305 | 110.44 | <0.001 |
| SumDbh | 0.6072 | 53.031 | 121.963 | <0.001 |
| $F_{\text{♂}}$, plot N4-HIGH | Adjusted $R^2=0.181$; p-value<0.001 AIC<sub>ref</sub>=335.73 | | | |
| PMis | -0.45 | 37.64 | 340.50 | 0.010 |
| SumDbh | 0.42 | 28.28 | 338.84 | 0.026 |
| Stature | Dom : 0.31 | 117.71 | 352.18 | <0.001 |
|  | Cod : 0.86 |  |  |  |
|  | Suppr : -1.17 |  |  |  |

**Figure 2.**
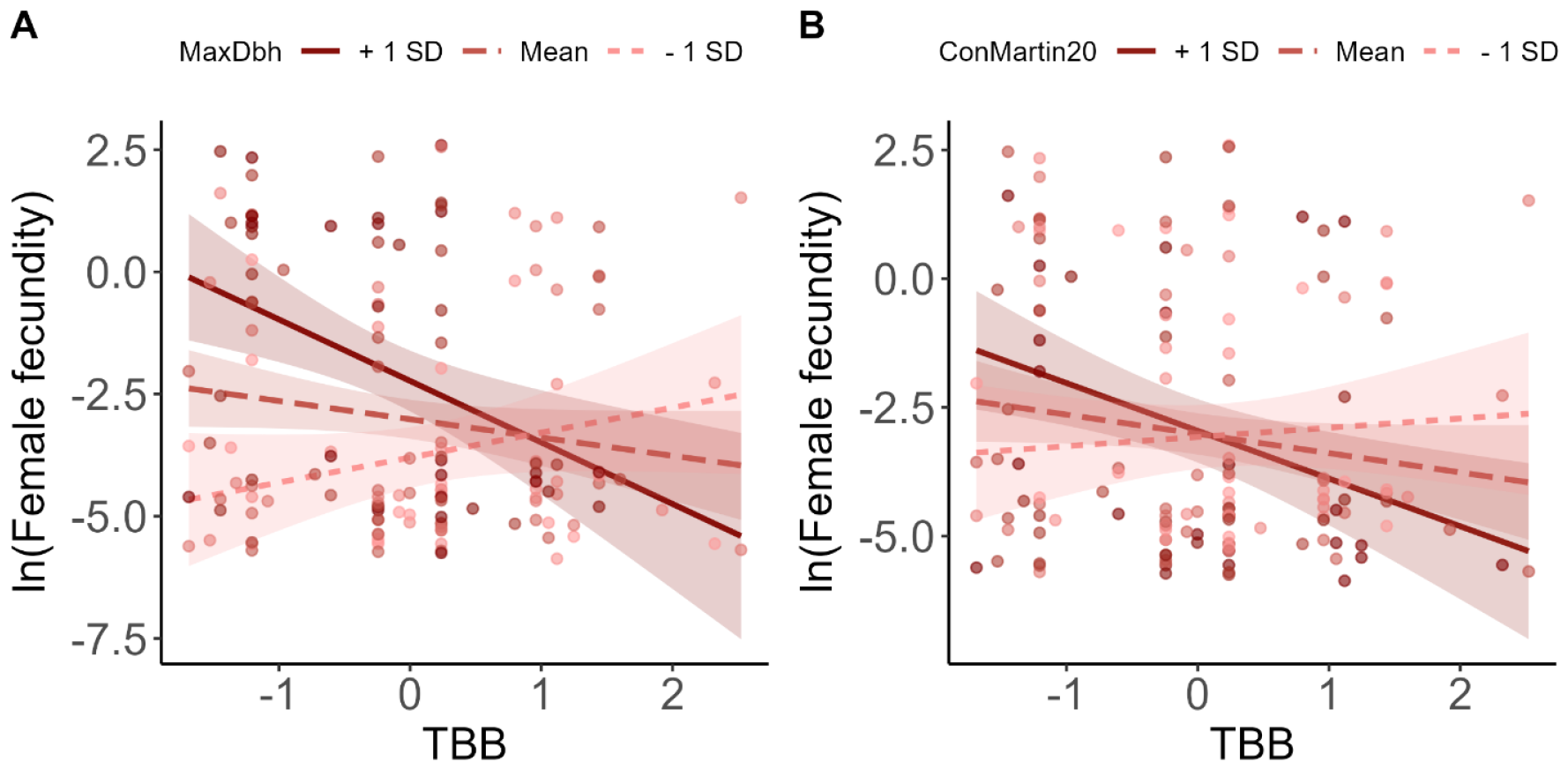
Interaction effects between TBB and size (A) or competition (B) on female fecundity at low altitude (plot N1-LOW). Predicted regression lines are plotted for three values of each moderator variable, corresponding to +/-1 standard deviation from the mean. Confidence interval at 95% are shown around each regression line. Dots are the observed values

The best model for female fecundity at plot N4-HIGH included TBB, size, and competition, as well as the interaction between TBB and size, although this interaction term was not significant. As the second-best model, without the competition term, performed nearly as well as the best one (ΔAICc = 0.56), we favored parsimony and kept it. Our results suggest that delayed TBB and increased competition may decrease female fecundity (β_TBB_=-0.34, p=0.052; β_ConMartin20_=-0.43, p=0.05), while female fecundity increased with tree size (β_MaxDbh_=0.78, p<0.001).

In contrast, earlier budburst only increased male fecundity at high altitude. The best model for male fecundity at plot N1-LOW showed a marked increase in male fecundity with tree size (β_SumDbh_=0.56, p<0.001), but no significant effect of TBB. At plot N4-HIGH (Fig. 3B), delayed TBB decreased male fecundity (β_TBB_=-0.41, p=0.019). Male fecundity also increased with tree size (β_SumDbh_=0.46, p=0.015), and depended on tree stature, with higher fecundity for codominant and dominant trees. Table S2 displays the results of all the fitted models of fecundity selection while Fig. S6 shows the residuals of the best models.

**Figure 3.**
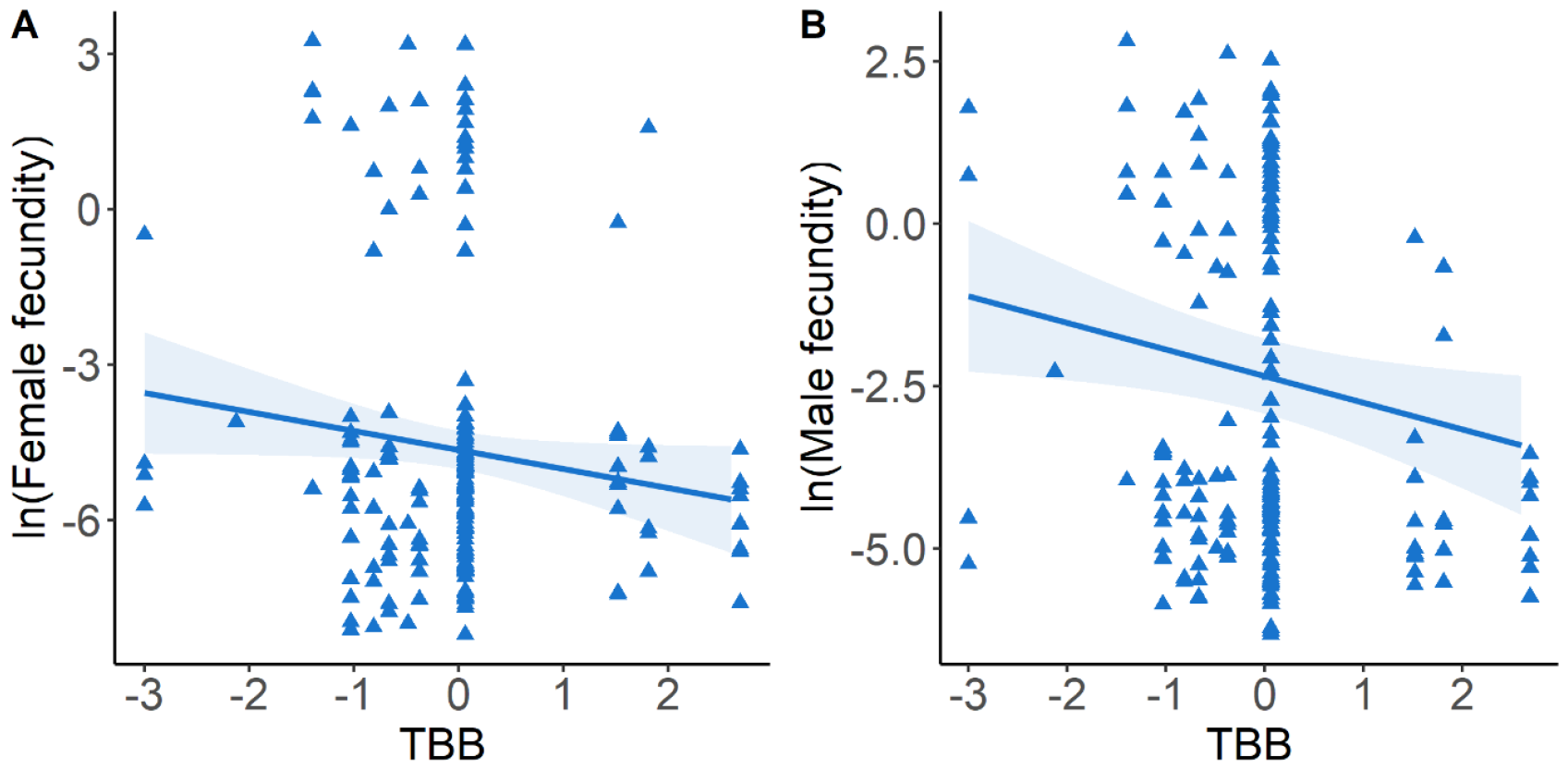
Relationship between TBB and female fecundity (A) or male fecundity (B) at high altitude (plot N4-HIGH). The lines are the predictions with their 95% confidence intervals, and triangles are the observed values.

### Estimation of assortative mating and phenological mismatch

Assortative mating, as estimated by the correlation in TBB between mating pairs, was significantly positive at plot N1-LOW (ρ=0.196, p<0.001) but not significant at plot N4-HIGH (ρ = −0.11, p = 0.09). These results were based on 713 seedlings assigned to a most-likely father: 397 among the 694 genotyped seedlings at plot N1-LOW (57%) and 316 among the 720 genotyped seedlings at plot N4-HIGH (44%). At plot N1-LOW, the joint distribution of phenological score (PSS) for parent pairs differed from the expected distribution under random mating (Fig. 4).

**Figure 4.**
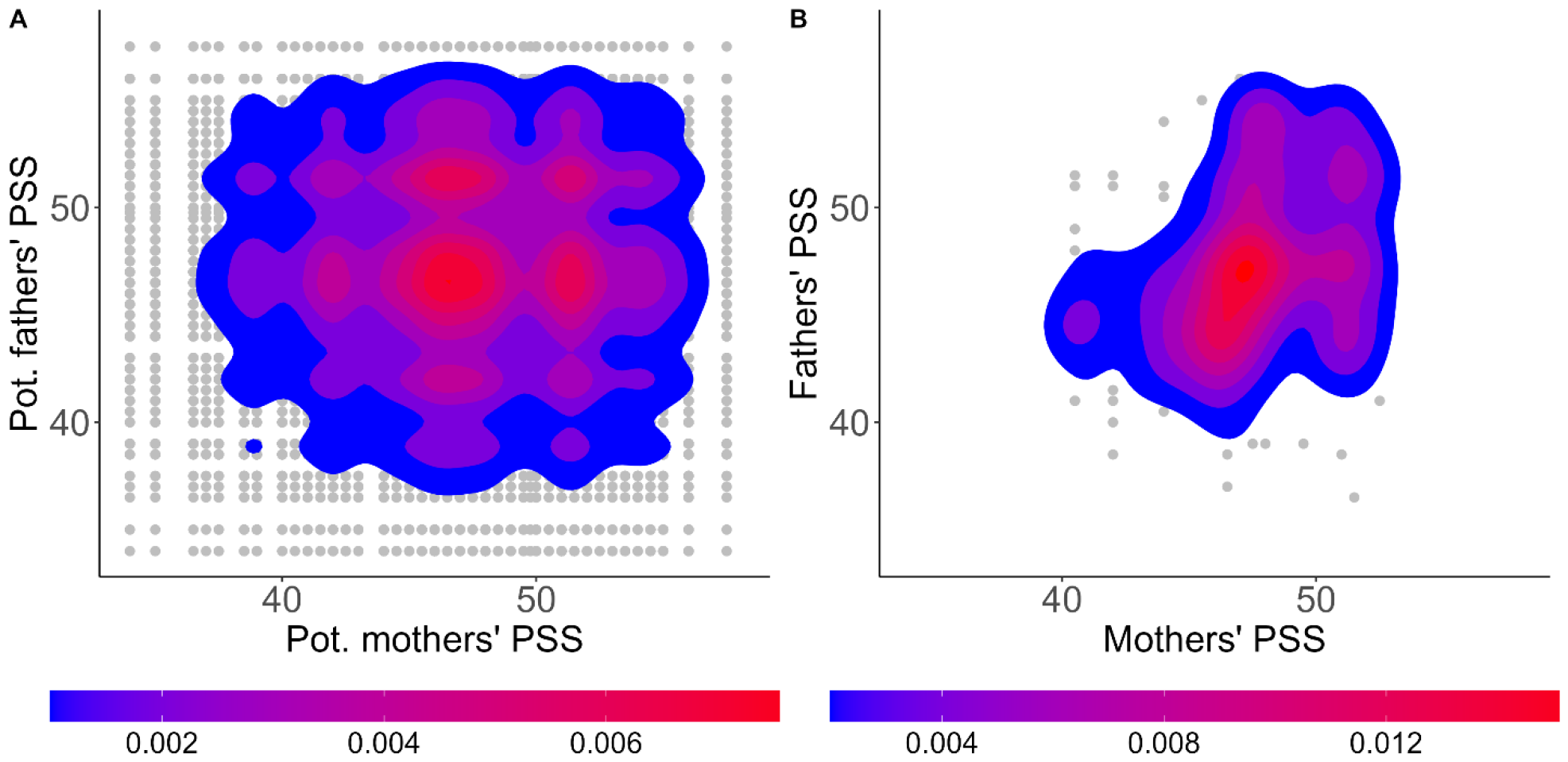
Join distribution of parent pairs’ phenological score (PSS) under random mating (A), and in realized mating events (B) at plot N1-LOW. A. The density of the data cloud was computed under the hypothesis that each tree mates once as male and once as female with all possible trees. B. Paternity analyses of seeds sampled on mother-tree allowed to identify mates’ pairs. See also Fig. S7.

For the analyses that follow, we selected |*PMis*|_*s*_ within a 20 m radius, referred to as PMis hereafter, as the most accurate estimator of phenological mismatch due to its high variation (Table S1). At plot N1-LOW, PMis decreased as phenological spread increased (corr_PMis-spread_ = −0.23, p-value=0.004). Conversely, the opposite trend was observed at plot N4-HIGH (corr_PMis-spread_ = 0.12, p-value=0.09, Fig. S8A). The relationship between TBB and PMis was found to be quadratic, with a TBB value that minimized PMis (Fig. S8B). This result is expected if TBB is not strongly spatially structured (Fig S9). The distributions of |*PMis*|_*s*_ and |*PMis*|_*m*_ at each plot can be seen in Figures S10 and S11.

### Sexual selection analyses for phenological mismatch, based on adult trees in situ

Only male fecundity variation was significantly affected by PMis (Fig. 5). In the best models, male fecundity decreased with increasing PMis at both plots N1-LOW (β_PMis_=-0.44, p<0.001) and N4-HIGH (β_PMis_=-0.45, p<0.001). Despite similar selection gradient values at both plots, the distribution of observed values of TBB and fecundity suggest stronger sexual selection at plot N1, in line with the stronger signal of assortative mating. Male fecundity increased with SumDbh at both plots, and for trees with codominant stature at plot N4. The results of all the fitted models of sexual selection are shown in Table S3, and the residuals of the best models are shown in Fig. S12.

**Figure 5.**
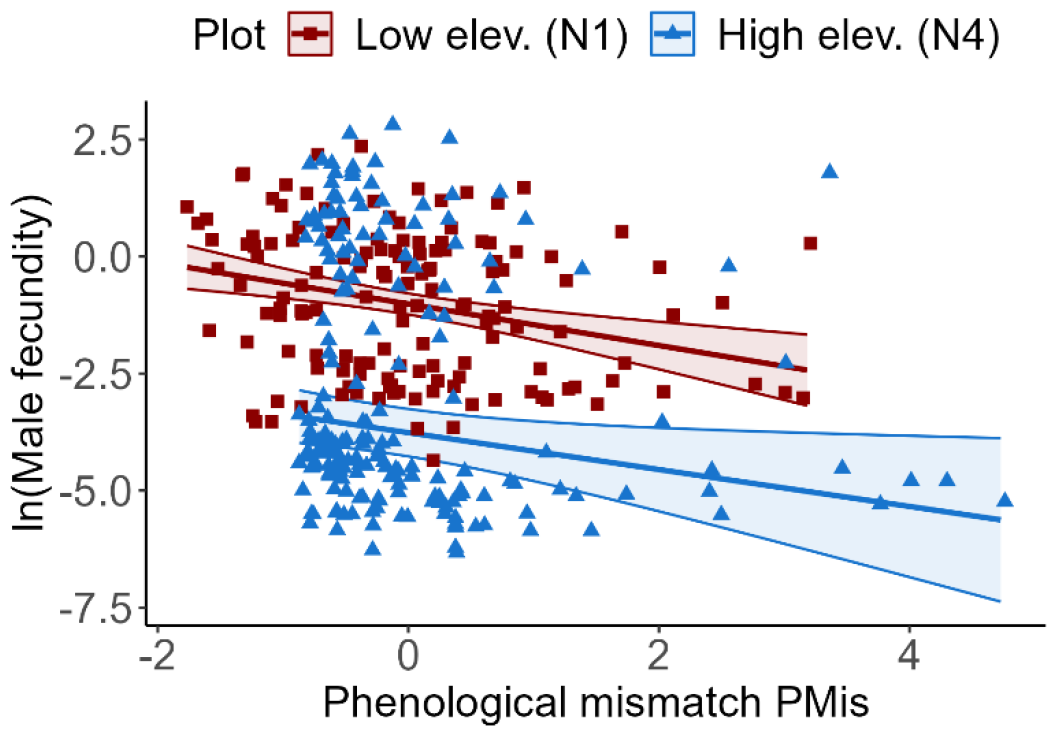
Relationship between phenological mismatch and male fecundity (Bateman’s gradient). The higher the phenological mismatch, the lower the opportunities for mating. The phenological mismatch, PMis, was estimated as the sum of absolute difference in TBB between a tree and each of its neighbors in a 20 m radius. Symbols represent observed values (square: plot N1-LOW; triangle: plot N4-HIGH) and lines are the prediction of the best sexual selection model.

We also tested compound best models, where both TBB and PMis were included as factors in the best model for fecundity selection. Only for male fecundity at plot N4-HIGH did the compound model outcompete the BestFec and BestSex models (Table S4). The effect of TBB in the compound model (β_TBB_=-0.34, p=0.045) was very similar to that of TBB in the BestFec model, indicating that sexual selection does not impact the estimate of fecundity selection. Regarding female fecundity, the effect of PMis was not significant in the compound model and the effect of TBB did not differ from that of the BestFec models, indicating that sexual selection does not impact the estimate of fecundity selection. The compound model did not show a significant effect of TBB on male fecundity at plot N1-LOW, and the effect of PMis was not different from that of the BestSex model.

### Estimation of stabilizing selection and standardized selection gradients on phenology

We found no evidence for stabilizing selection on TBB through a significant effect of TBB2 on either female or male fecundity. However, the significant effect of PMis on male fecundity in plots N1-LOW and N4-HIGH illustrates a form of stabilizing selection on TBB, as lower PMis are obtained for average TBB due to the quadratic relationship between PMis and TBB (Figure S8B).

Standardized selection gradients (Table S5) suggest that selection for earlier TBB through female fecundity is slightly higher on plot N4-HIGH (β_TBB_’=-0.43) than on plot N1-LOW (β_TBB_’=-0.24), although these differences are not significant due to large standard errors. Selection for earlier TBB through male fecundity at plot N4-HIGH was in the same order of magnitude (β_TBB_’=-0.30) as through female fecundity. The directional selection for reduced phenological mismatch with neighbors was slightly higher on plot N1-LOW (β_|PM|_’=-0.40) than on plot N4-HIGH (β_|PM|_’=-0.16), although these differences are also not significant.

### Growth selection analyses for TBB, based seedlings in the common garden

Growth selection analyses revealed a significant effect of TBB on seedling growth (Table 5): both diameter and height growth significantly decreased with delayed budburst (increasing TBB). In addition, growth increased with increasing initial size, and growth was reduced for seedlings originating from plot N4-HIGH compared to those from plot N1-LOW. As expected, the variance in growth was significantly structured by block and family (Table S6). A more detailed analysis showed the expected strong negative effect of water stress on growth. Moreover, the negative effect of delayed budburst on growth (although much lower than that of treatment) was higher in the water stress treatment (Table S7).

**Table 5:** Selection on seedling growth in diameter (Dgrowth) and height (Hgrowth). Selection on phenology was assessed through the effect of TBB on seedling growth, accounting for the effects of plot (N1-LOW or N4-HIGH), initial size (initD or initH), and common garden design (with Block and Family included as random effects). The global significance of each fixed term was assessed based on the Sum Of Squares (SoSq) and the F-value (F-test), while the effect of TBB within each plot was assessed based on the t-value (Student test). See Table S7 for a more complex model including treatment.

A- Diameter growth
| Term | npar | SoSq | F-value | p-value | Effect | St. error | t-value | p-value |
| --- | --- | --- | --- | --- | --- | --- | --- | --- |
| Plot | 1 | 2.86 | 4.12 | 0.042 | -0.345 | 0.387 | -0.89 |  |
| initD | 1 | 228.76 | 330.18 | <0.001 | 0.291 | 0.016 | 18.66 |  |
| Plot:TBB11 | 2 | 15.80 | 11.40 | <0.001 | N1: -0.015<br>N4: -0.021 | 0.006<br>0.005 | -2.80<br>-3.94 | 0.005<br><0.001 |

B- Height growth
| Term | npar | SoSq | F-value | p-value | Effect | St. error | t-value | p-value |
| --- | --- | --- | --- | --- | --- | --- | --- | --- |
| Plot | 1 | 67549 | 11.83 | 0.001 | -76.940 | 35.256 | -2.18 |  |
| initH | 1 | 17753 | 3.11 | 0.078 | 0.046 | 0.020 | 2.33 |  |
| Plot :TBB11 | 2 | 53894 | 4.72 | 0.009 | N1: -0.188<br>N4: -1.545 | 0.511<br>0.504 | -0.37<br>-3.07 | 0.712<br>0.002 |

## Discussion

In this study, we estimated fecundity, sexual and viability selection by combining field and common garden data with parentage analyses in order to better understand the selection regime on spring phenology in European beech. Our main results were that fecundity selection on female fitness and viability selection on seedlings growth both favor early phenology, while sexual selection on male fitness through assortative mating modulates this trend (stabilizing selection). Furthermore, this study confirmed that environmental variation (here, altitudinal differences) can also have a major impact on the potential for contemporary evolution. Overall, this study highlights the value of in situ phenotypic selection analyses to better understand the evolutionary potential of tree populations (Bontemps et al. 2017; Alexandre et al. 2020; Westergreen et al. 2023).

### Earlier budburst increases female fecundity and seedling growth, with contrasted effects of drought stress

Our findings that earlier budburst increased female fecundity of adult trees in situ and seedlings growth in the common garden is consistent with the pervasive phenotypic selection for early phenology documented in plants (Geber and Griffen 2003; Munguía-Rosas et al. 2011; Austen and Weis 2015). These findings also contradict the expectation of stabilizing selection on vegetative and reproductive phenology, that would be driven by the balance between the benefits of avoiding frost damages on the one hand and maximizing the duration of the growing season on the other hand. Hence, we seem to face a similar paradox to the one observed for flowering phenology in short-lived plants, and for which Austen et al. (2017) already proposed four explanations: (1) selection through other fitness components may counter observed fecundity selection for early flowering; (2) asymmetry in the flowering-time–fitness function may make selection for later flowering hard to detect; (3) flowering time and fitness maybe condition-dependent; and (4) selection on flowering duration is largely unaccounted for. Before detailing how this study sheds ligth on mechanisms related to the first explanation (see the second part of this discussion), we can first add a fifth possible explanation to this list, related to temporally fluctuating selection in long-lived plants. Indeed, since we estimated selection during a single reproductive episode, we can not exclude the possibility that different patterns of selection may be observed in different years due to year-specific climatic conditions. A review has already suggested that changes in selection direction across years are common in vertebrates (Siepielski, Dibattista, & Carlson, 2009). In our case in particular, we may not have had favorable conditions to observe selection for later budburst driven by late frosts, as they did not occur in the year when we sampled seeds and seedlings. Selection for later budburst through late frosts can be expected to be a strong selective force(Westergren et al. 2023), because late frost damage can severely reduce the photosynthetic capacity of adult trees and thus their seed development and maturation and/or reduce seedling survival. However, this selection may occur only occasionally in the balance of other selections that occur each year with a more moderate intensity. Finally, it should be noted that among the possible sources of selection not generally considered in classical phenotypic selection analyses are those related to interspecific interactions. For example, in this multispecies ecosystem located at the ecological boundary between Mediterranean and mountainous climates, interspecific competition could be involved in shaping patterns of selection, reinforcing the tendency for beech to flush earlier than competing species (Palacio-Lopez et al., 2020).

However, we found some evidence against the general pattern of fecundity selection for earlier TBB through female fecundity. At low altitude, directional fecundity selection for early budburst was found only for the larger trees, while the smaller trees tended to show the opposite pattern, or at least, no fecundity gain with early budburst. Larger trees also had an earlier budburst, resulting in a consistent signal of directional selection for TBB and for this size category. Such contrasting selection gradients on TBB among neighboring trees suggest that different ecological strategies exist within the same drought-prone population, likely due to some microenvironmental heterogeneity. This pattern can be related to the “growth-stress survival” trade-off (Grime 1977; Grubb 1998), where slower development (small trees) and delayed budburst can be viewed as a drought tolerance strategy. This is consistent with a previous study on the same low-altitude plot (Bontemps et al. 2017), where trees displaying late budburst were also associated with small size, low leaf water content and other traits (e.g., high leaf mass per area) symptomatic of a water-saving strategy; at the same time, trees displaying early budburst were associated with large size, high leaf water content and other traits (e.g. low water use efficiency) symptomatic of a water-uptake strategy. In contrast with this “growth-stress survival” trade-off for adult trees facing different levels of water stress in situ, we found the opposite trend in the common garden, where the positive relationship between early budburst and higher seedling growth was slightly stronger in the drought-stress treatment of the experiment. The latter results can be interpreted as the fact that an early budburst allows seedlings to start photosynthesis when the conditions are the most optimal for growth (i.e., before drought) and can be viewed as a “drought-escape” strategy. Taken together, our results suggest that the patterns of selection on phenology may vary across life history stages (reviewed in Schluter et al. 1991). For example, Vitasse (2013) had previously showed that the earlier ontogenic stage of seedlings in the understory explains their earlier leaf emergence. Here, we suggest that the adaptive response to drought may differ between young and mature trees.

### Mating opportunities limit male fecundity, and drive stabilizing selection on TBB

Another key finding of this study, consistent with the first explanation proposed by Austen et al. (2017), is that stabilizing selection on male mating success through assortative mating can modulate fecundity selection for earlier phenology. To begin, this study is among the first ones to demonstrate and estimate assortative mating on spring phenology in a tree species. Furthermore, consistent with Bateman’s principle, we found that increasing phenological mismatch with neighbors, as a proxy for decreasing mating opportunities, affected male but not female fecundity. Thus, phenological variation among trees within a stand creates opportunities for sexual selection, and may drive stabilizing selection on TBB through the male function. Such stabilizing selection has already been observed in a pollen-limited population of *Quercus lobata*, where trees that flowered early or late set fewer acorns than trees that flowered at the peak of the population (Koenig et al., 2012). Our study generalizes this result to cases where pollen does not limit fruit set.

Assortative mating has important evolutionary and ecological consequences (Jiang et al. 2013), and assortative mating for phenological traits in particular may strongly influence the evolutionary response to climate change (Godineau, Ronce, & Devaux, 2021; Soularue & Kremer, 2014; Whittet et al., 2017). However, the standard measure of assortative mating based on the observation of individual synchrony of flowering schedules (Weis et al. 2005, 2014) is hardly applicable to forest trees. Therefore, the potential assortative mating for phenological traits has mostly been investigated between tree populations, by measuring the differences in the timing of pollen shedding between oak populations along temperature clines (Whittet et al. 2017) or by inferring the latitudinal origin of pollen in open-pollinated pine progeny grown in common gardens (Nilsson 1995). Another common approach is to estimate mating system parameters using genetic markers; such studies have proposed assortative (or disassortative) mating as a general mechanism leading to higher (or lower) relatedness between mated individuals than expected by chance (Hardy et al., 2019; Ismail & Kokko, 2020; Monthe et al., 2017). Here, we applied the approach widely used in animal species to quantify assortative mating (e.g., Jiang et al. 2013): first, we used paternity analyses to infer mated pairs *a posteriori*, and second, we computed the correlation of spring phenology among members of mated pairs. To our knowledge, this is one of the few studies to assess effective assortative mating for spring phenology in a tree species, by combining budburst phenology data with marker-based paternity analyses (see also Gérard et al. 2006; Lagache et al. 2014; Larue et al. 2022). Our approach showed significant assortative mating for spring phenology at the lower plot, where budburst spread over 17 days. The correlation in vegetative phenology between mating pairs was moderate (ρ=0.19) as compared to the range reported in the literature (e.g. 0.05–0.63 within the same old-field community, Weis et al. 2014). In the upper plot, the more rapid development of leaf unfolding, which occurred over only 13 days, may explain why assortative mating was not detected, although we cannot rule out that other factors limit mating, such as higher canopy density at higher elevations.

As we expected due to assortative mating, we found that flowering timing synchronized with close neighbors maximizes mating success through the male function, but does not significantly affect the female fecundity. Indirectly, this favors intermediate timing of bud burst since in the absence of a strong spatial structure, males with intermediate TBB are those most synchronized with their neighbors. To our knowledge, this study is the first to test and validate Bateman’s principle in a tree, likely due to the difficulty of estimating the number of mates in these species that produce a large number of offspring. We used the phenological mismatch as a proxy for (potential) mating opportunities rather than the mating success, which could have been estimated based on paternity analyses (Tonnabel et al. 2019), because our sampling design, of only 35.3 seeds per mother tree, may underestimate the contribution of rare fathers. The effect of phenological mismatch could be related to stabilizing selection on TBB, as the phenological mismatch is a quadratic function of TBB. However, and surprisingly, we did not find the expected consequence of a significant quadratic relationship between effective fecundity and TBB. This could be due to different abilities to detect significant linear coefficients (from the slope of the regression line) as compared to quadratic coefficients (from the curvature of the fitness surface).

The observed effect of mating opportunities on MEMM estimates of fecundity is counterintuitive, as these estimates are claimed to be effective estimates of basic fecundity (Oddou-Muratorio et al. 2018). This is likely because the effect of the phenological mismatch is not included in the MEMM model we used; thus, any effect of phenological mismatch on individual reproductive success is incorporated into the estimate of individual fecundity. In the same way that MEMM models the effect of the relative positions of putative parents and offspring on fecundity through the pollen dispersal kernel (spatial assortative mating), we could also model the effect of phenological mismatch on fecundity in MEMM (temporal assortative mating, as done in Gérard et al. 2006; Gleiser et al. 2018; Larue et al. 2022). In this case, the estimated fecundity would no longer depend on the mating opportunities. This option would be interesting to include in future developments of MEMM.

### Altitudinal variation of selection on spring phenology and overall evolutionary potential of the studied beech population

Although selection gradients for each component of selection (female fecundity, male fecundity, sexual selection on male fecundity) did not differ significantly among altitudes, this study highlighted a number of qualitative indications that selection for earlier phenology (i.e., precocity) is stronger overall at high altitude than at low altitude in the population studied. First, selection for precocity through female fecundity was reinforced by selection for precocity through male fecundity only at high altitude. Second, selection for precocity through female fecundity was modulated by the interaction effect between size and TBB only at low altitude. Third, assortative mating, the fuel for sexual stabilizing selection through male mating success, was stronger at low altitude. Stronger selection for earlier phenology at high latitude is consistent with the physiological expectation that the length of the growing season strongly constrains the level of resources acquired through photosynthesis (Keenan et al. 2014; Richardson et al. 2006). It is also consistent with the simulation study of Gauzere et al. (2020a) showing that selection for earlier budburst is stronger under conditions that are more limiting to reproductive development, i.e., in cold environments.

On a quantitative point of view, the standardized directional selection gradients on spring phenology estimated in this study (β’) ranged between −0.43 and −0.24. This suggests a rather strong magnitude of selection, using the meta-analysis of Kingsolver et al. (2001) as a reference (where a mean |β’|-value of 0.22 was found across all traits, with a median |β’|-value of 0.08 for life-history/phenological traits). This metanalysis also reported higher value of |β’| for selection via fecundity or mating success (median |β’| = 0.18) than for selection via survival (median |β’| = 0.09), supporting the strong directional selection on fecundity estimated here. Considering the high level of narrow-sense heritability estimated for phenological score sum in the population at low altitude (h2=0.84–0.92; Bontemps et al. 2016), our results may indicate a high evolutionary potential for spring phenology in the studied population. Such strong selection gradients are likely to reflect strong selective pressures on phenology that may constraint population demographic growth in both cold and warm environments. This supports the hypothesis that phenology is an important determinant of survival and fecundity, consistent with studies that use it to predict the distribution range of plant species (Chuine & Beaubien 2001, Gauzere et al. 2020a). However, the high evolutionary potential of spring phenology measured in the studied beech population does not guarantee by itself its ability to adapt to the multiple effects of ongoing climate change. In particular, there is increasing evidence that emerging drought stress is causing massive mortality even in areas previously spared by drought (Hartmann et al. 2022). Whether the genetic response of spring phenology to increased summer temperature combined with extreme drougth stress will allow beech populations to adapt is difficult to predict without a dedicated predictive modelling approach (e.g., Oddou-Muratorio & Davi 2014). However our results show that accounting for genetic differences in phenological schedules and their ecological significance can greatly improve scenarios of future population adaptation to drought and late frost stress.

Spring phenology has been defined as a “magic trait”, because it simultaneously affects fitness through its influence on growing season (and thus survival and fecundity) and contributes to nonrandom mating (Servedio et al., 2011; Soularue & Kremer, 2014). Previous simulation studies have shown how environmental variation can cause populations to diverge for a selectively neutral trait that causes assortative mating (Kirkpatrick 2000; Soularue and Kremer 2012). Consequently, some patterns of clinal genetic variation in phenological traits observed in forest trees may be generated solely by the effects of assortative mating and gene flow, in the absence of divergent selection. When divergent selection and assortative mating for TBB occur simultaneously, Soularue and Kremer (2014) predicted that genetic clines can be either inflated or constrained by assortative mating, depending on species life history. Finally, a recent study predicted the evolution of either suboptimal plasticity (reaction norms with a slope shallower than optimal) or hyperplasticity (slopes steeper than optimal) for TBB in the presence of assortative mating, whereas optimal plasticity would evolve under random mating (Soularue et al. 2022). These different simulation studies considered prescribed, single-trait models of divergent selection, in which a single optimal value maximizes fitness within each population. Given the intertwined effects of sexual, fecundity and viability selection on phenology and the variation in fitness landscapes for budburst along temperature and drought gradients shown in this study, we suggest that future eco-evolutionary models of phenological shifts should integrate these features in a mechanistic and multidisciplinary framework (Donohue et al. 2015, Lamarins et al. 2022). Such an approach could allow quantitative assessment of which type of selection (viability, fecundity, sexual selection) currently dominates the selection regime on spring phenology, and evaluate whether the genetic response to these different types of selection will allow beech populations to adapt to ongoing climate change.

## Acknowledgements

We thank Olivier Gilg, Frank Rei, Norbert Turion, Frédéric Jean and Mehdi Pringarbe (INRAE UEFM) for sample collection, field measurements, and seed germination. We thank Anne Roig and Matthieu Lingrand (INRAE URFM) for genotyping and managing genetic databases. We also thank Hendrik Davi, Jeanne Tonnabel and Ophélie Ronce for discussion on previous version of this manuscript.

## Data, scripts, code, and supplementary information availability

Data (two files: dataAdultField.txt and dataGrowthCommonGarden.tab), Scripts of statistical analyses (two files: SelectionAnalyses_adult.html and GrowthSelectionAnalysis_seedlings.html) and Supplementary figures and tables (one file: SuplMaterial_V10.pdf) are available online: https://doi.org/10.57745/ZVPNXX

## Conflict of interest disclosure

The authors declare that they comply with the PCI rule of having no financial conflicts of interest in relation to the content of the article. S.O.M and E.K. are recommenders of PCIEvolBiol.

## Funding

The study was funded by the EU ERA-NET BiodivERsA projects TIPTREE (BiodivERsA2-2012-15), the bilateral ANR project EXPANDTREE (2013 - SVSE 7) and the ANR project MeCC (ANR-13-ADAP-0006). This study was supported by two PhD thesis allocation grants awarded to: (1) A.B. by Région PACA and by ECODIV research division at INRAE; (2) J.G. by ECODIV and MIA research divisions and CLIMAE metaprogram at INRAE.

## Authors contributions

Research conceptualization: S.O.M, A.B., E.K., J.G. Design or development of research methods and tools: S.O.M, E.K., A.B., J.G. Data collection, analysis, management, interpretation: S.O.M, A.B., E.K., J.G. Validation: S.O.M, E.K. Manuscript writing: S.O.M. (first draft), E.K., J.G., A.B.

## Notes

https://doi.org/10.57745/ZVPNXX

### Competing Interest Statement

The authors have declared no competing interest.

### Summary of Updates

This article has now been peer-reviewed and recommended by PCIEvolBiol

