## Supplementary Tables and Figures for "Interplay between fecundity, sexual and growth selection on the spring phenology of European beech (*Fagus sylvatica* L.)"

**Supplementary online Figures and Tables for the manuscript: “Interplay between fecundity, sexual and growth selection on the spring phenology of European beech (Fagus sylvatica L.)”, by S. Oddou-Muratorio, A. Bontemps, J. Gauzere, and E.K. Klein**

**Figure S1: Spring mean (top panels) and minimal (bottom panels) temperatures in year 2009 at plots N1-LOW (left side) and N4-HIGH (right side).**

We obtained the daily temperatures from 1959 to 2015 from the closest grid point of the SAFRAN database, and elevation effects on temperature were simulated using linear models and meteorological data recorded from 2007 to 2013 in to the two studied populations. The main thick line represents the simulated monthly average (top panels) or monthly minimal (bottom line panels) temperatures at the focal year (2009), while the grey area represents the simulated 95% variation interval from 1959 to 2015, and the two dotted lines are the minimal and maximal values over this period. The stars represent the monthly average (top panels) or monthly minimal (bottom panels) temperatures recorded in to the two studied populations at year 2010.

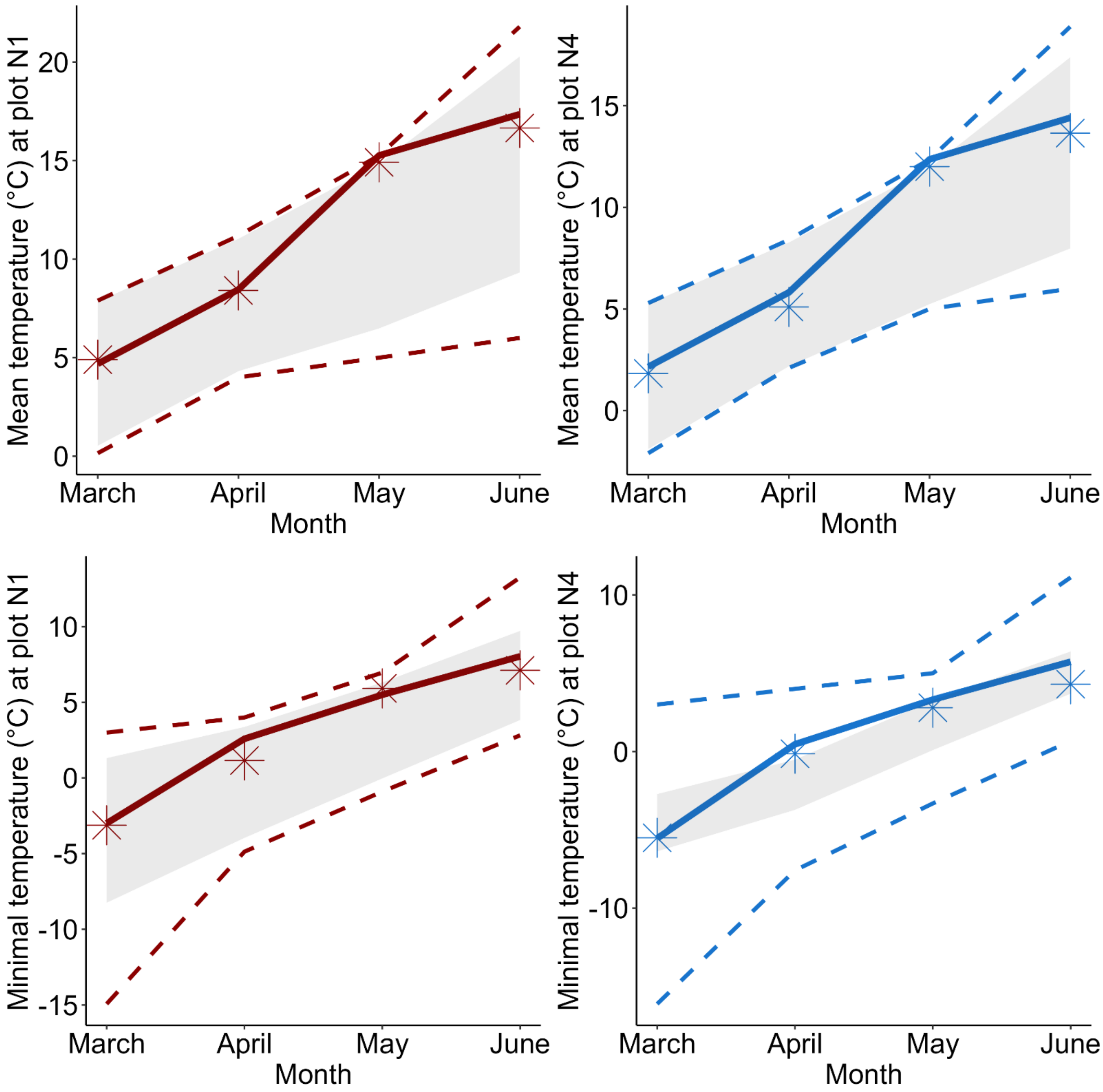

**Figure S2: Standard operating procedure for measuring budburst phenology.**

Five stages were used to follow the bud burst dynamics (from 1 to 5). The pictures below allow to match these five stages to closest BBCH reference stage, considering the BBCH scale adapted to trees and shrubs (https://tempo.pheno.fr/Presentation/Variables-mesurees).

| Stage 1: buds are dormant (BBCH 0) | Stage 4: the leaves are emerging (BBCH 08-09) |
| --- | --- |
| 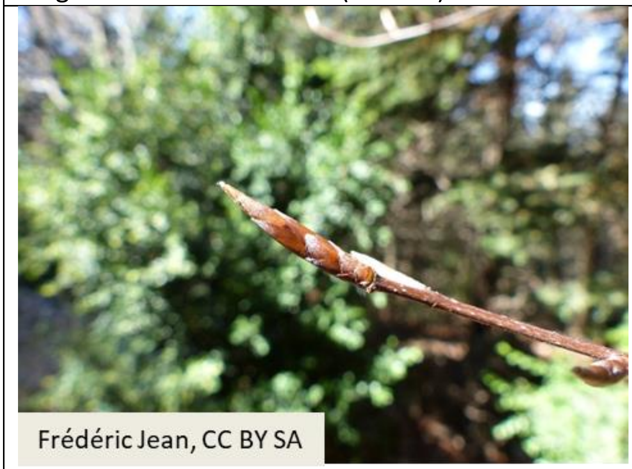  | 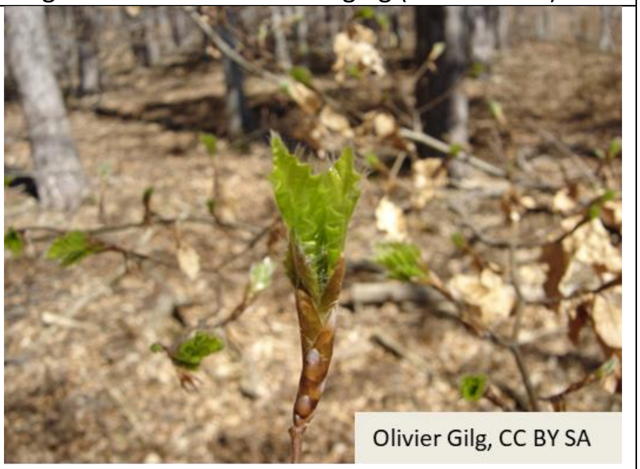  |
| Stage 2: buds are swelling (BBCH 1) | Stage 5: 90% of leaves are spread out (BBCH 19) |
| 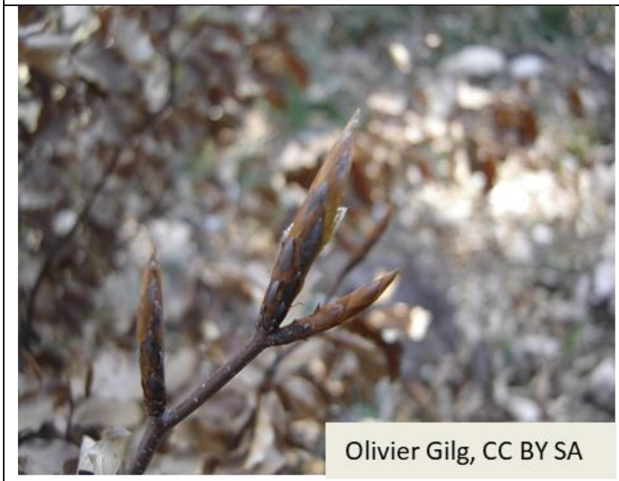  | 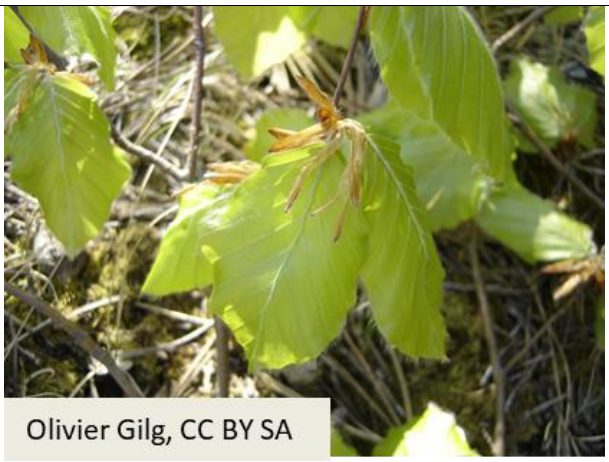  |
| Stage 3: bud scales are broken<br>(BBCH 07) | |
| 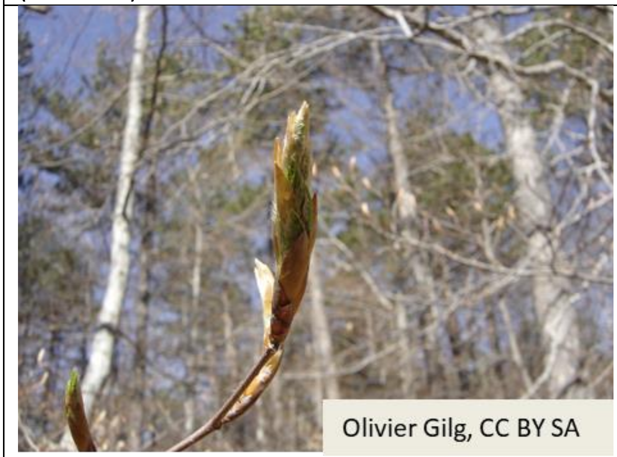  | |

**Figure S3. Variation of the timing of budburst (TBB).**

A. Histogram of TBB (in julian days) in both stands. Dashed lines represent the mean TBB value within each stand (N1-LOW: 20th of April; N4-HIGH: 4th of May) B. Relationship between TBB and circumference. The correlation was significantly negative at plot N1-LOW ($\rho_{TBB\text{-}circ}$ = -0.15, p-value=0.007) but not at plot N4-HIGH ($\rho_{TBB\text{-}circ}$ = -0.02, p-value=0.12).

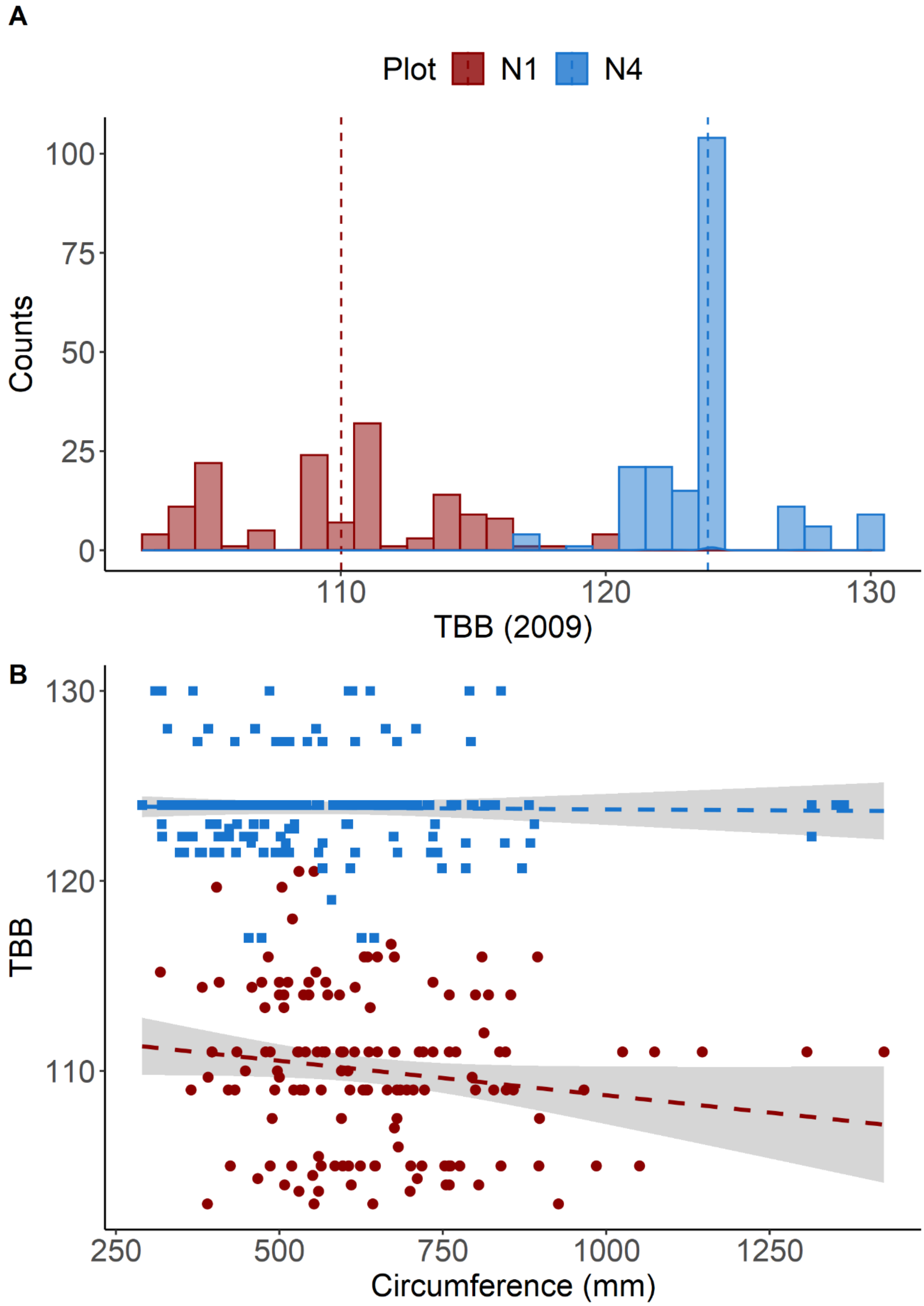

**Figure S4. Variation of phenological spread.**

A. Histogram of spread in both stands. Dashed lines represent the mean spread value within each stand (N1-LOW: 4.8 days; N4-HIGH:2.9 days). B. Relationship between TBB and spread. The correlation was significantly positive at plot N1-LOW ($\rho_{TBB\text{-}spread}$ = 0.57, p-value<0.01) but not at plot N4-HIGH ($\rho_{TBB\text{-}spread}$ = 0.09, p-value=0.20).

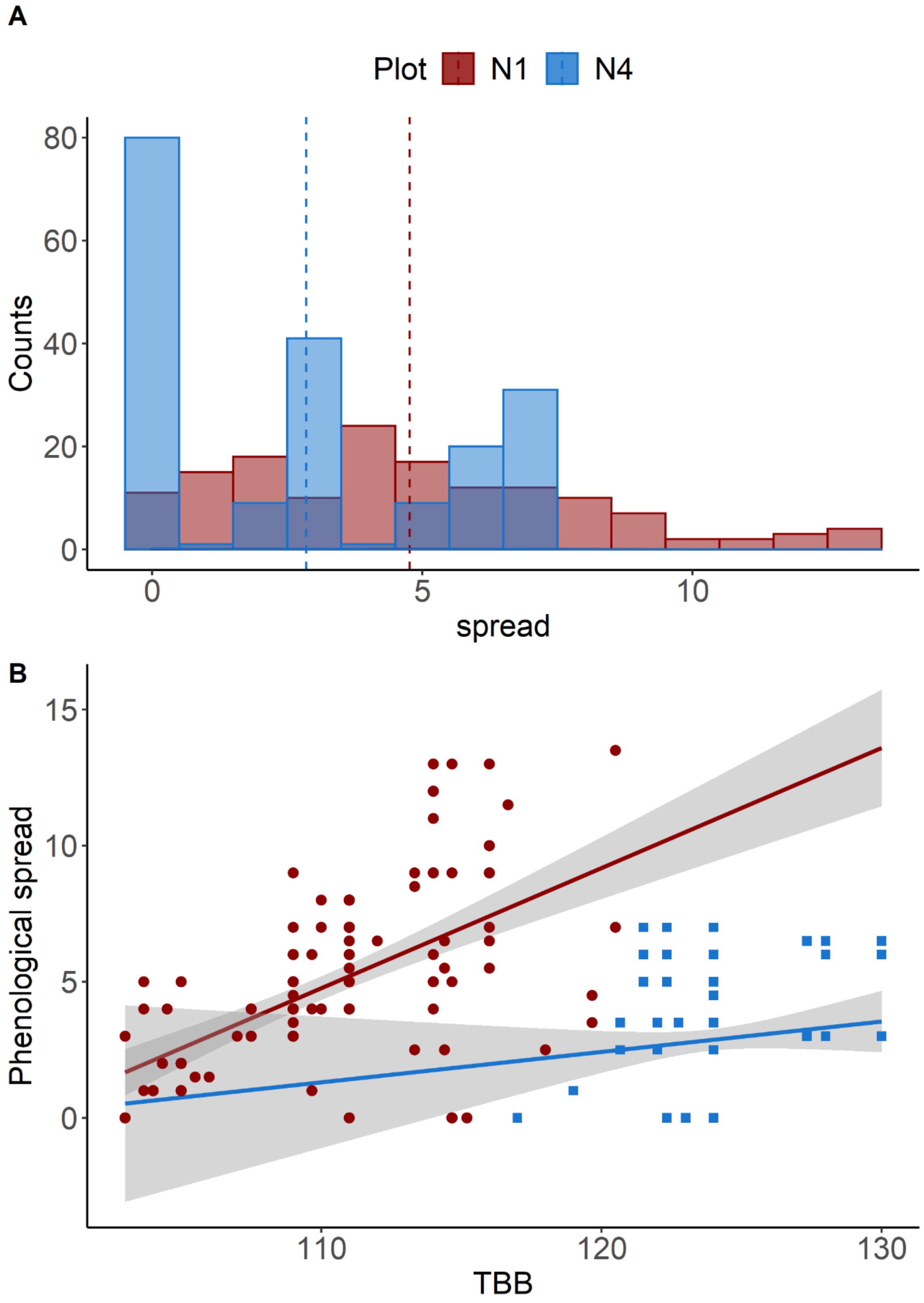

**Figure S5: Distribution of individual (A) male and (B) female fecundities at both plots N1-LOW and N4-HIGH**.

**A.** Male fecundities were estimated using MEMM based on maternal progenies collected on 20 mother-trees per plot. The logscale representation below illustrates the bimodality of the male fecundities (MF) distribution, with a peak on the left of the dashed line of negligible fecundities (i.e., not significantly different from the initial value).

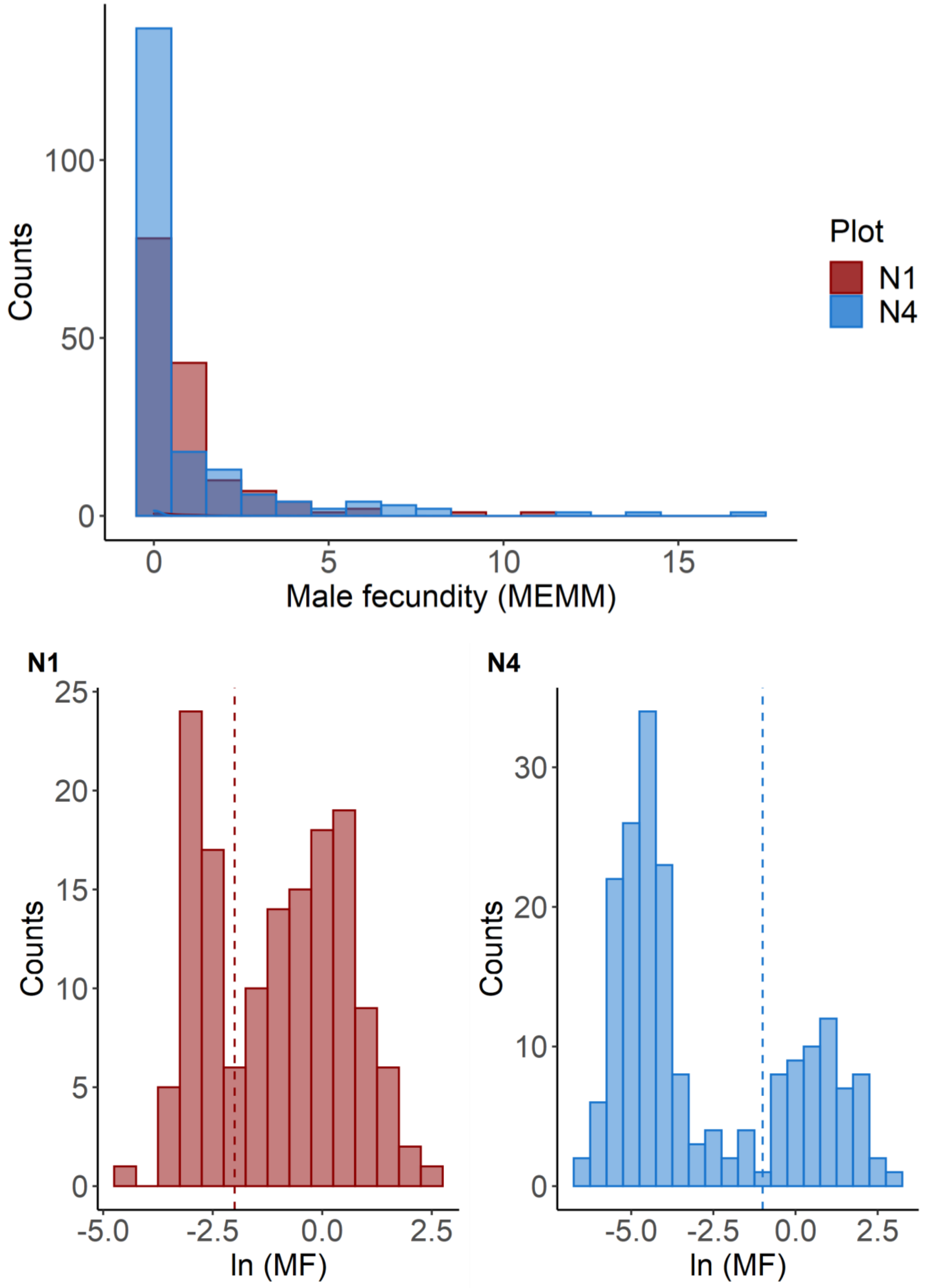

**B.** Female fecundities were estimated using MEMMseedlings based on established seedlings. The logscale representation below illustrates the bimodality of the female fecundities (FF) distribution, with a peak on the left of the dashed line of negligible fecundities (i.e., not significantly different from the initial value).

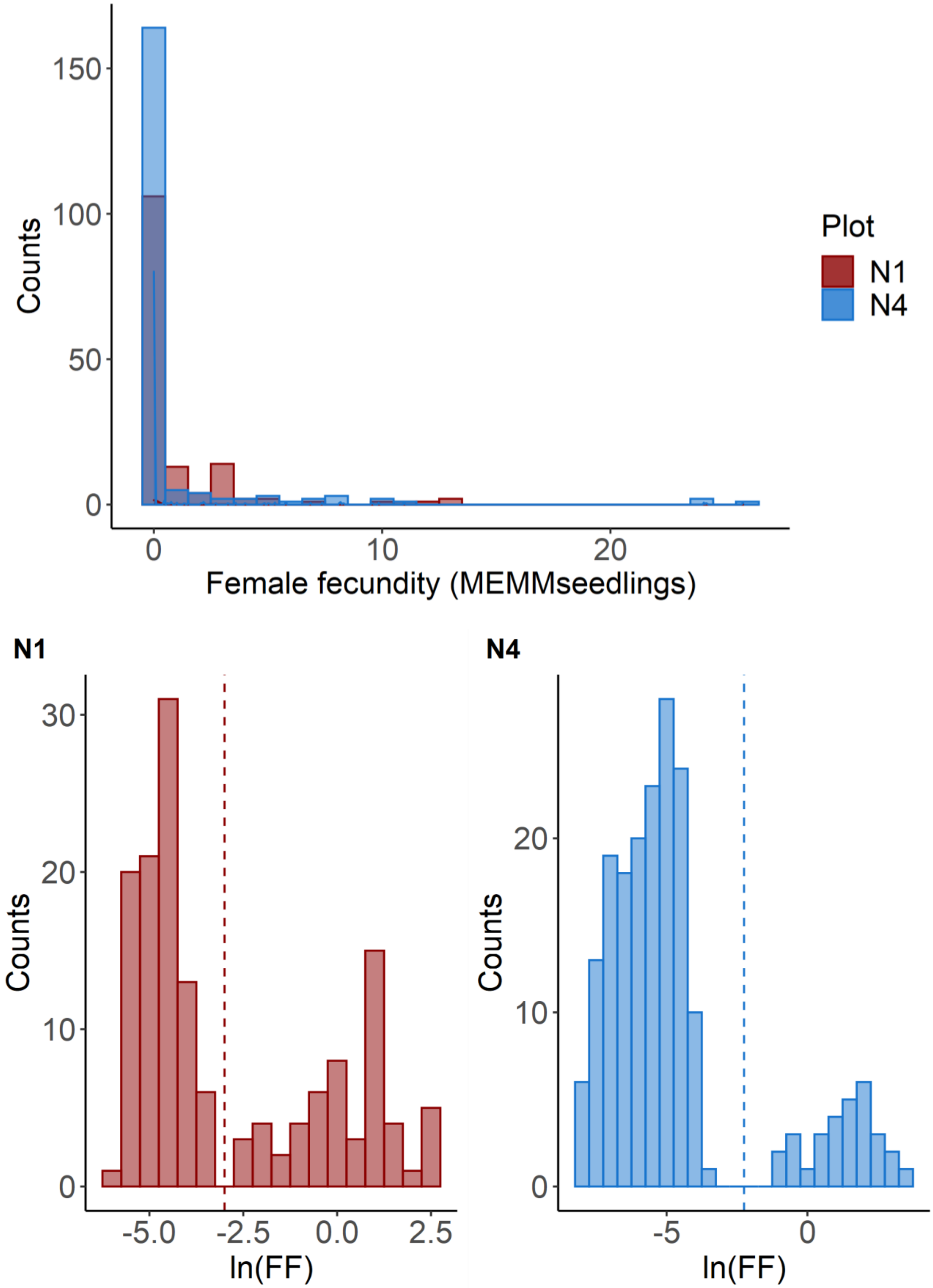

**Figure S6 Quality control of the fit of linear models for the estimation of fecundity selection.**

*(A) Fecundity selection model on female fecundity at plot N1-LOW (model M7.fec)*

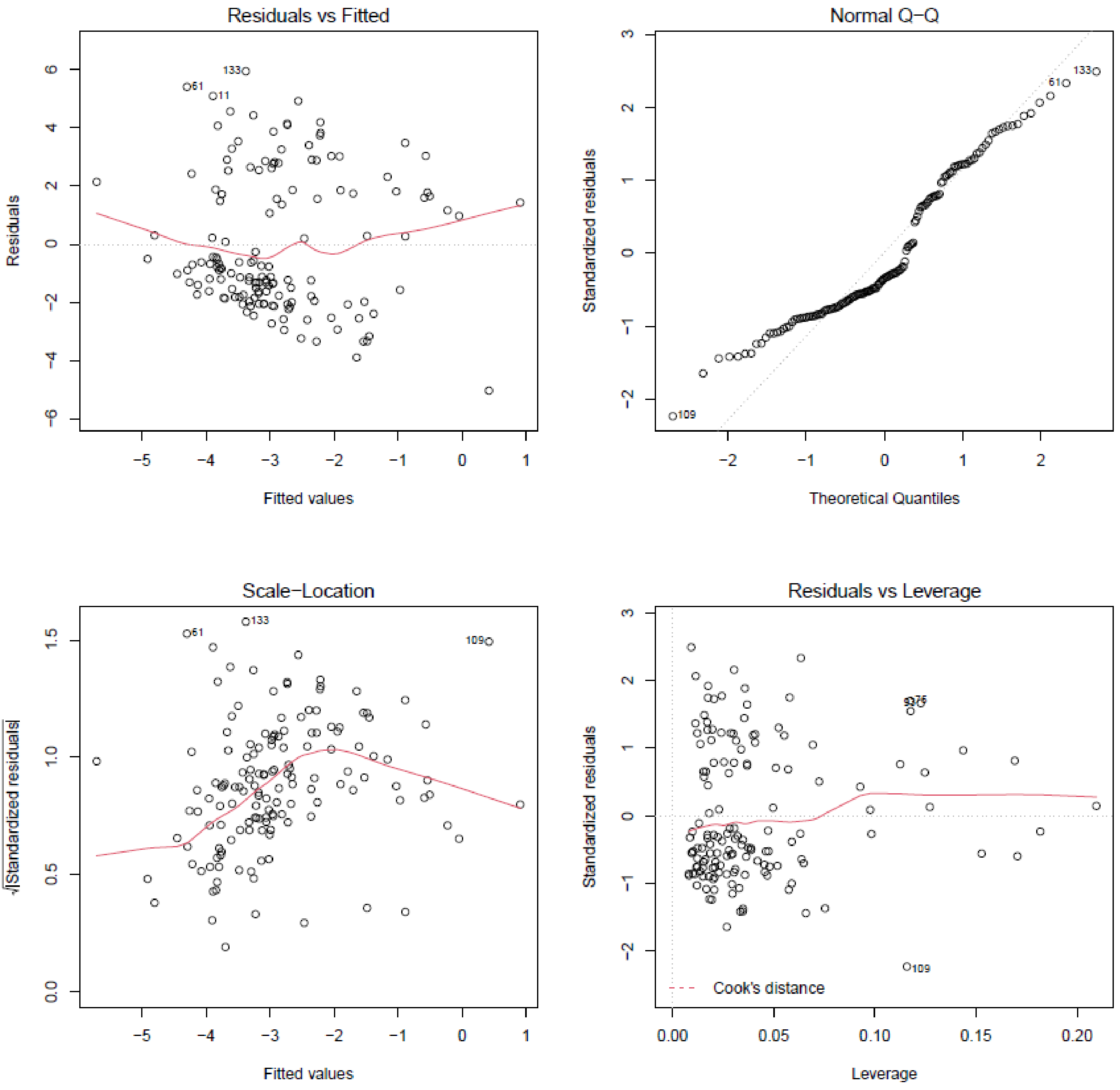

*(B) Fecundity selection model on female fecundity at plot N4-HIGH (model M5.fec)*

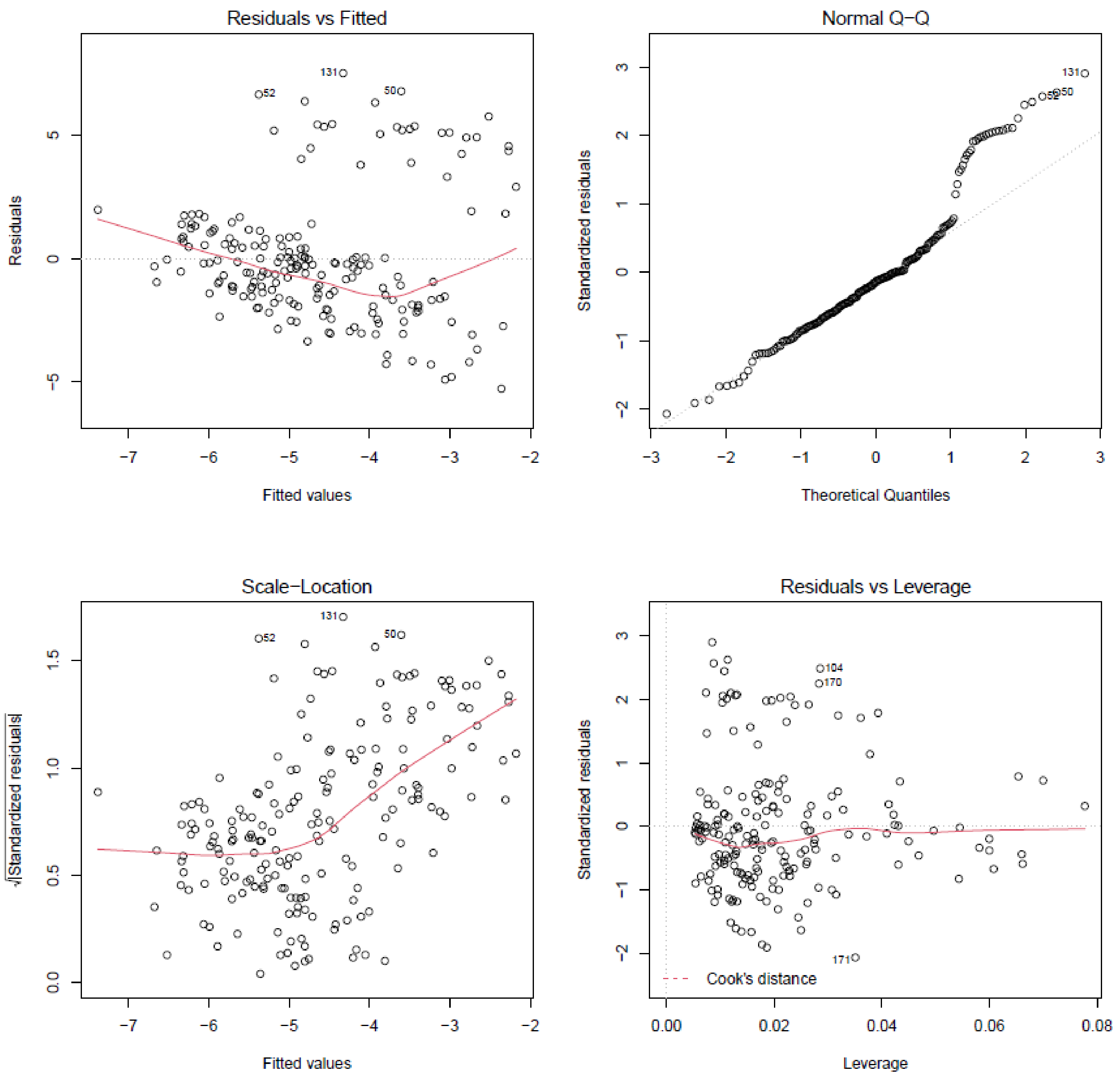

*(C) Fecundity selection model on male fecundity at plot N1-LOW (model M2.fec)*

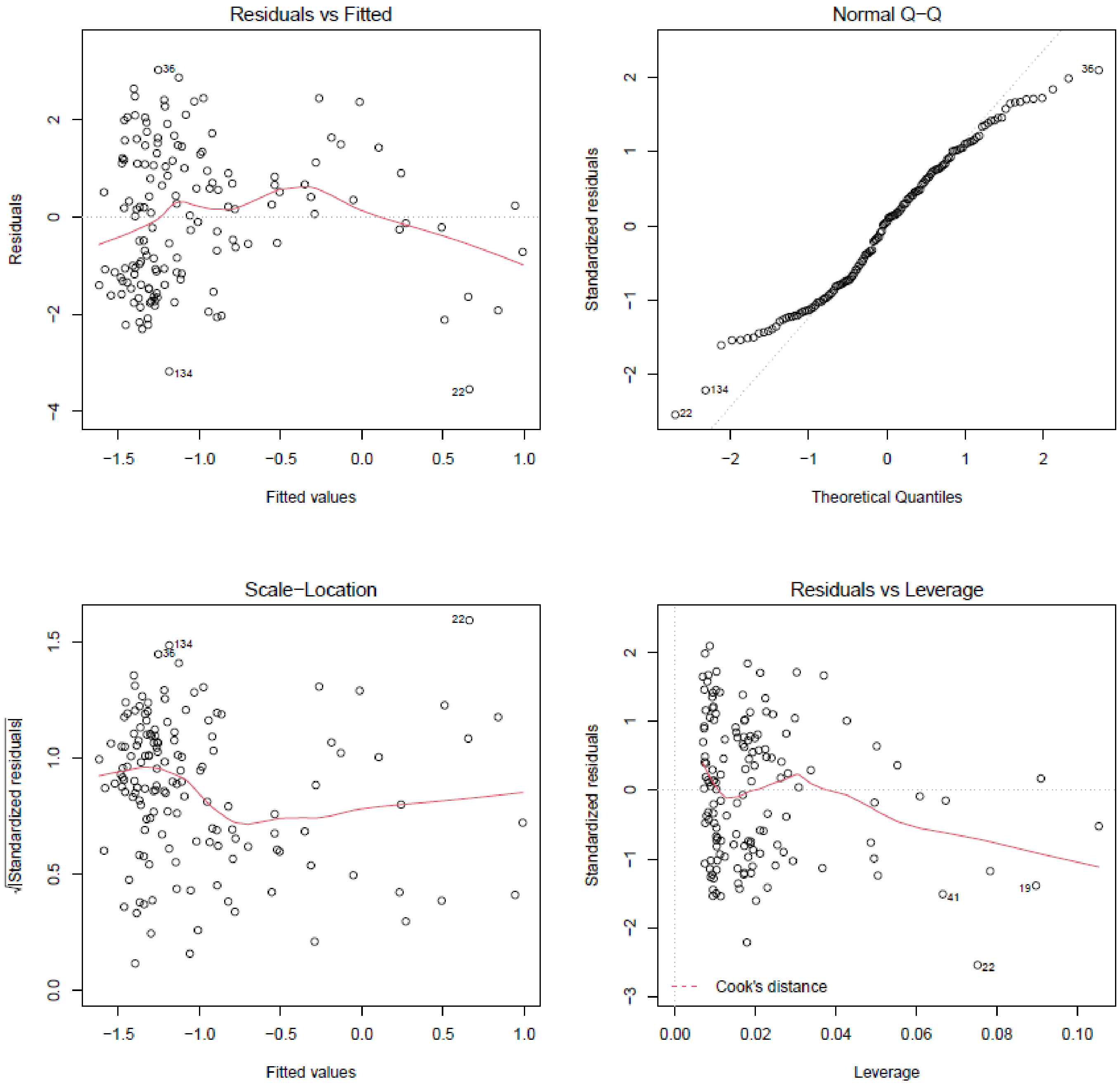

*(D) Fecundity selection model on male fecundity at plot N4-HIGH (model M4.fec)*

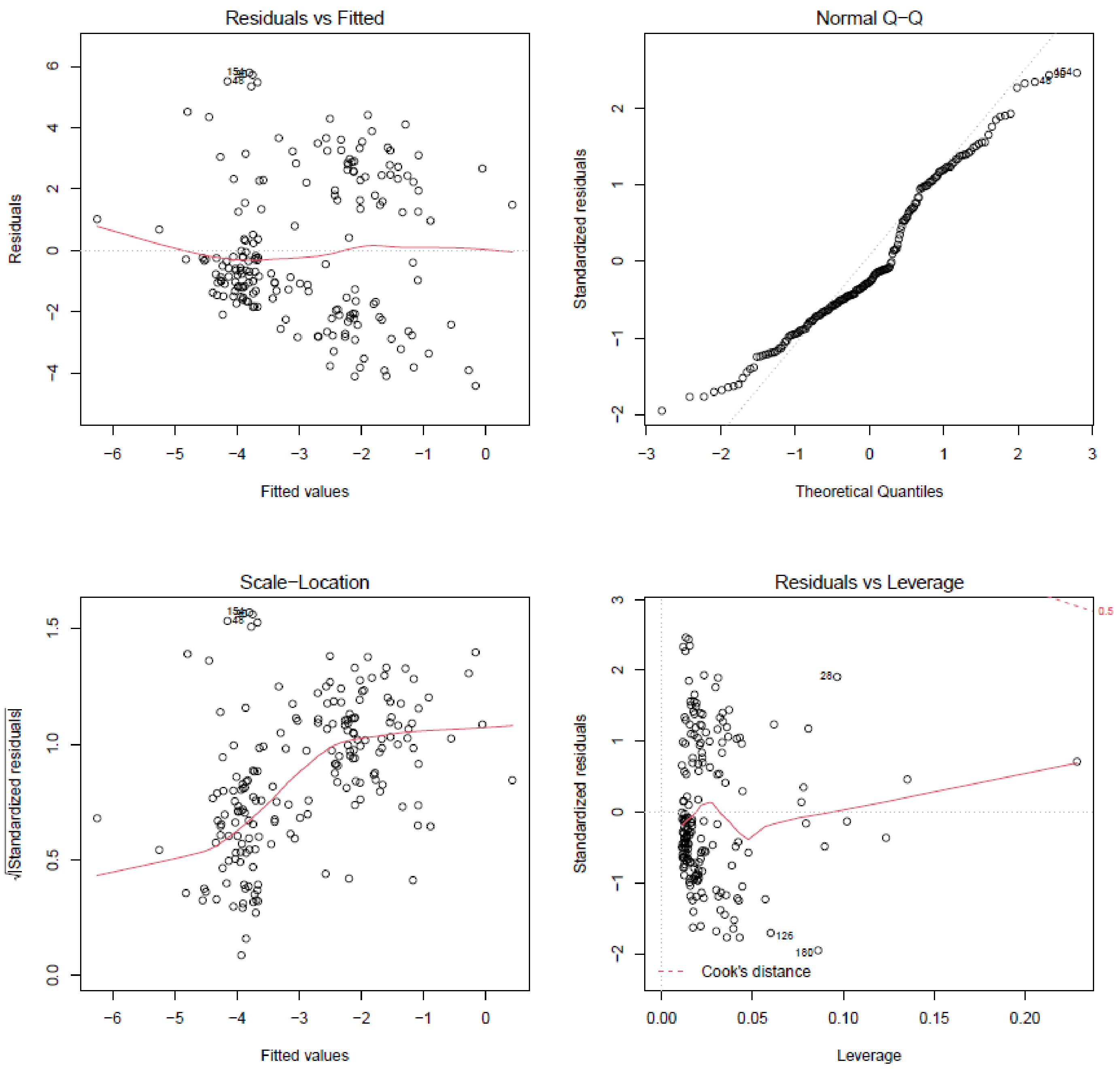

**Figure S7: Join distribution of parent pairs' TBB under random mating (A), and in realized mating events (B) at plot N1-LOW.**

A. The density of the data cloud was computed under the hypotheses that each tree mated in turn as male and female with all possible trees. B. Paternity analyses of seeds sampled on mother-tree allowed to identify mates' pairs.

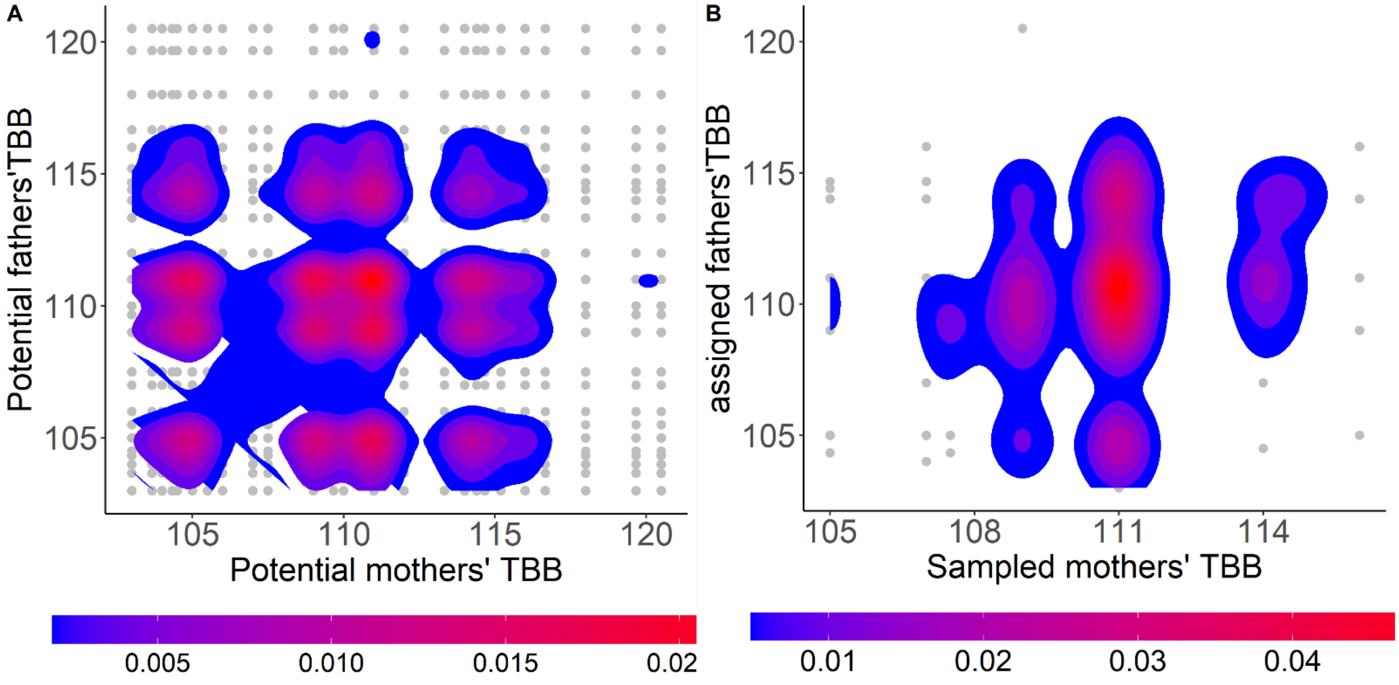

**Figure S8 Relationship between phenological mismatch (PMis) and A. phenological spread; B. the timing of budburst (TBB) at both plots.**

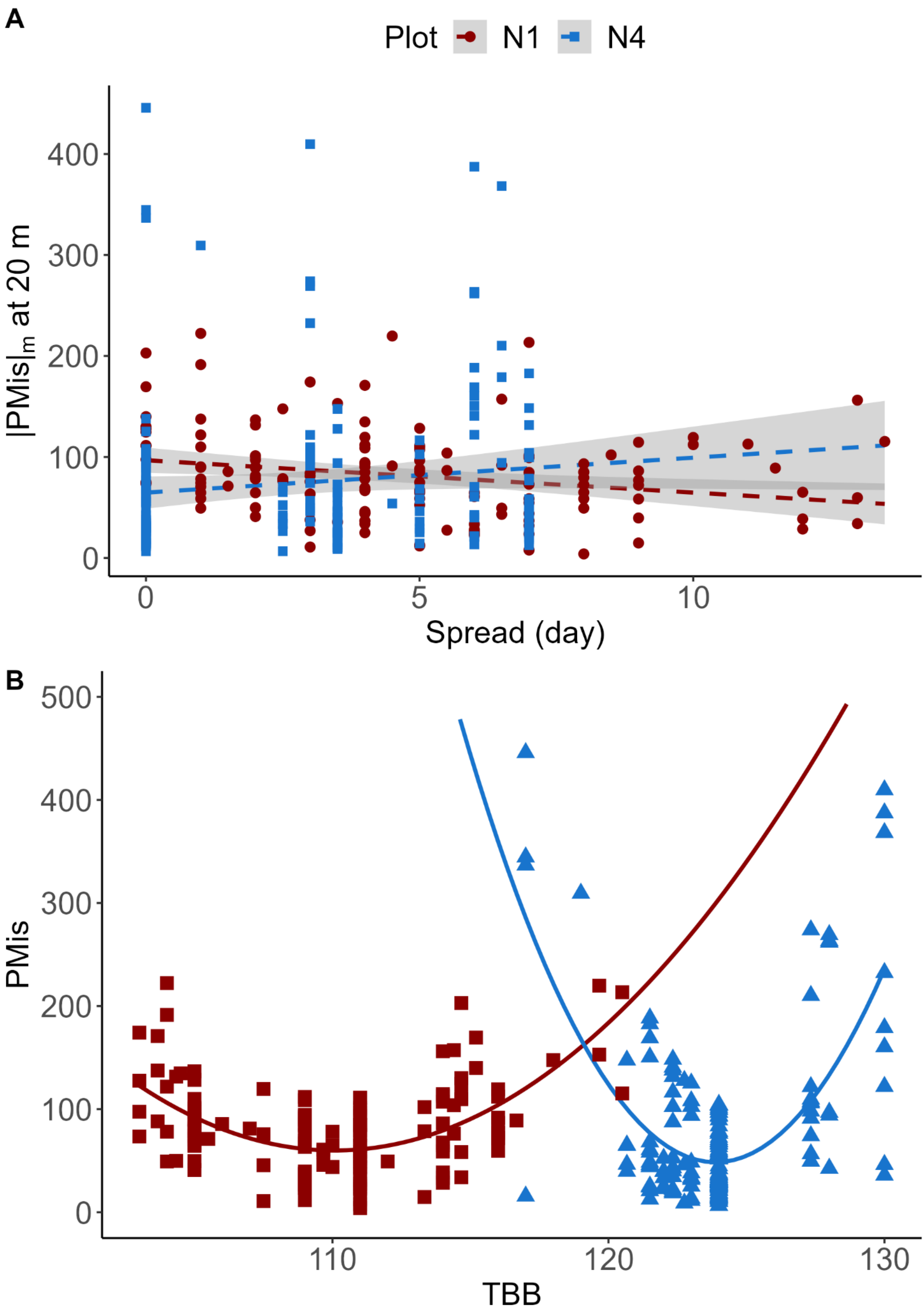

**Figure S9 Spatial autocorrelation of TBB at both plots (top: N1-LOW; bottom: N4-HIGH), as depicted by the variation of Moran's index I among pairs of individuals.**

At plot N1-LOW, the regression of Moran's I index against logarithm of distance (indicative of spatial autocorrelation) was significant only for male fecundity. At plot N4-HIGH, the regression of Moran's I index against logarithm of distance was significant only for female fecundity

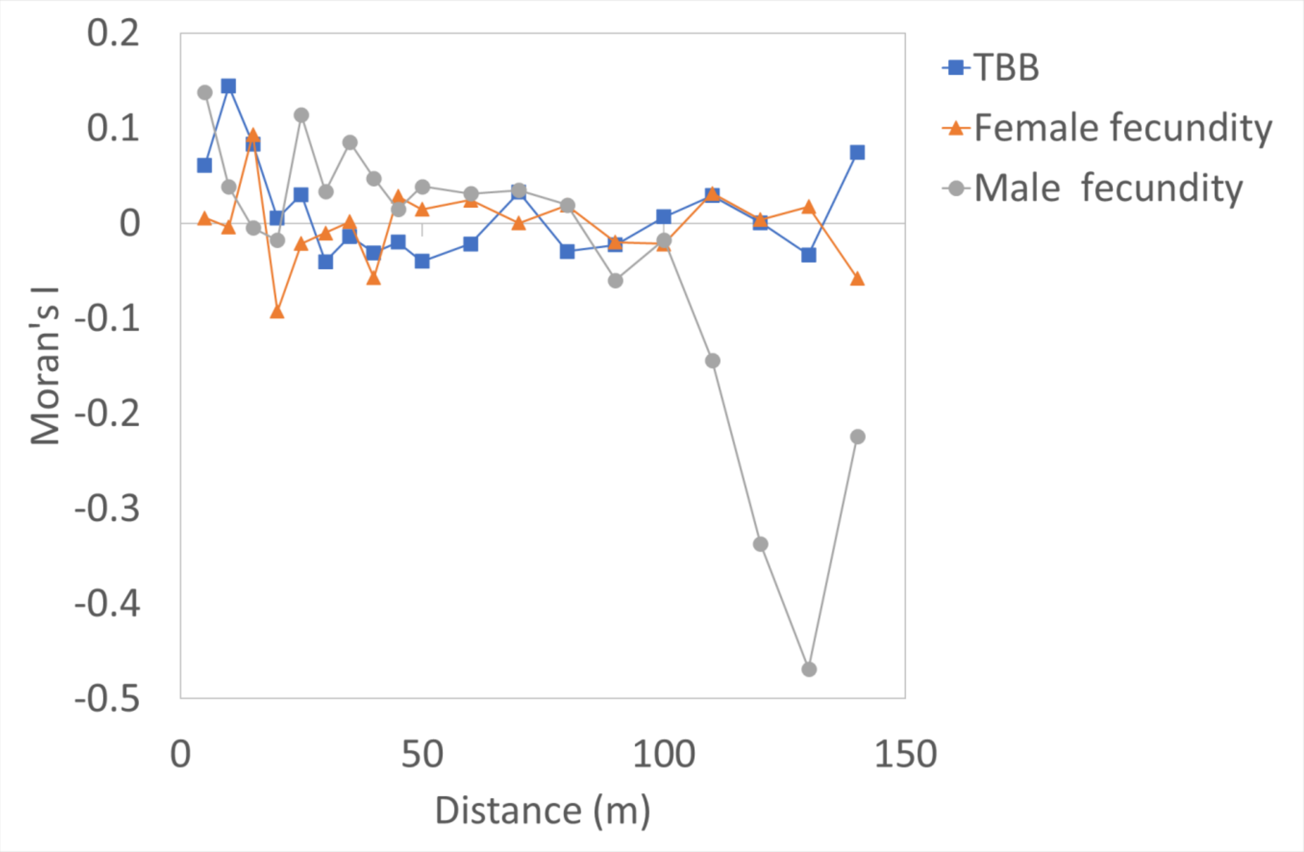

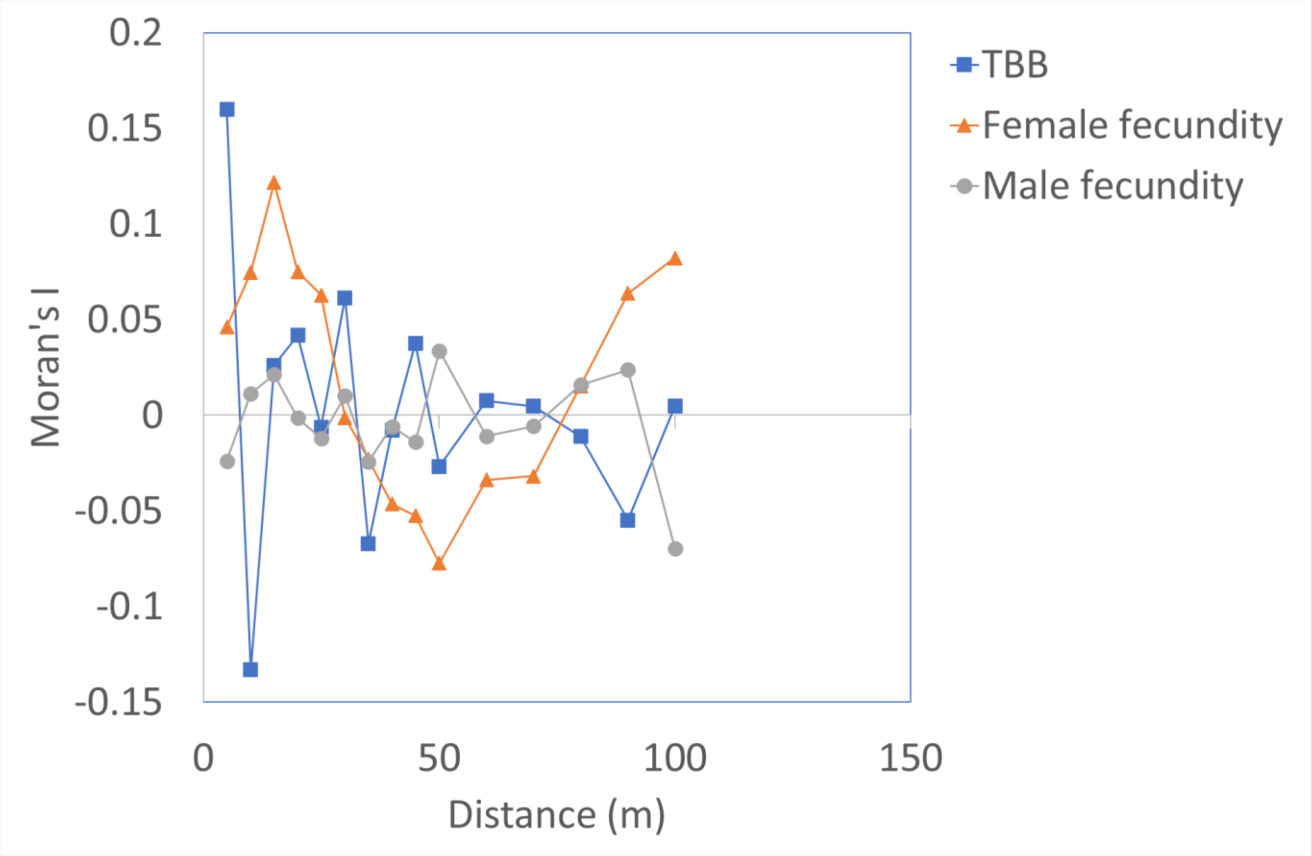

**Figure S10. Distribution of phenological mismatch (|PMis|$_s$).**

|PMis|$_s$ was computed as the sum of absolute differences in TBB between a focal tree and its neighbors in various radius at plots N1-LOW (red bars) and N4-HIGH (blue bars). The dashed lines represent mean values in each plot, illustrating that |Pmis|$_s$ was always higher at low that at high altitude. See also the variance of |Pmis|$_s$ in Table S1.

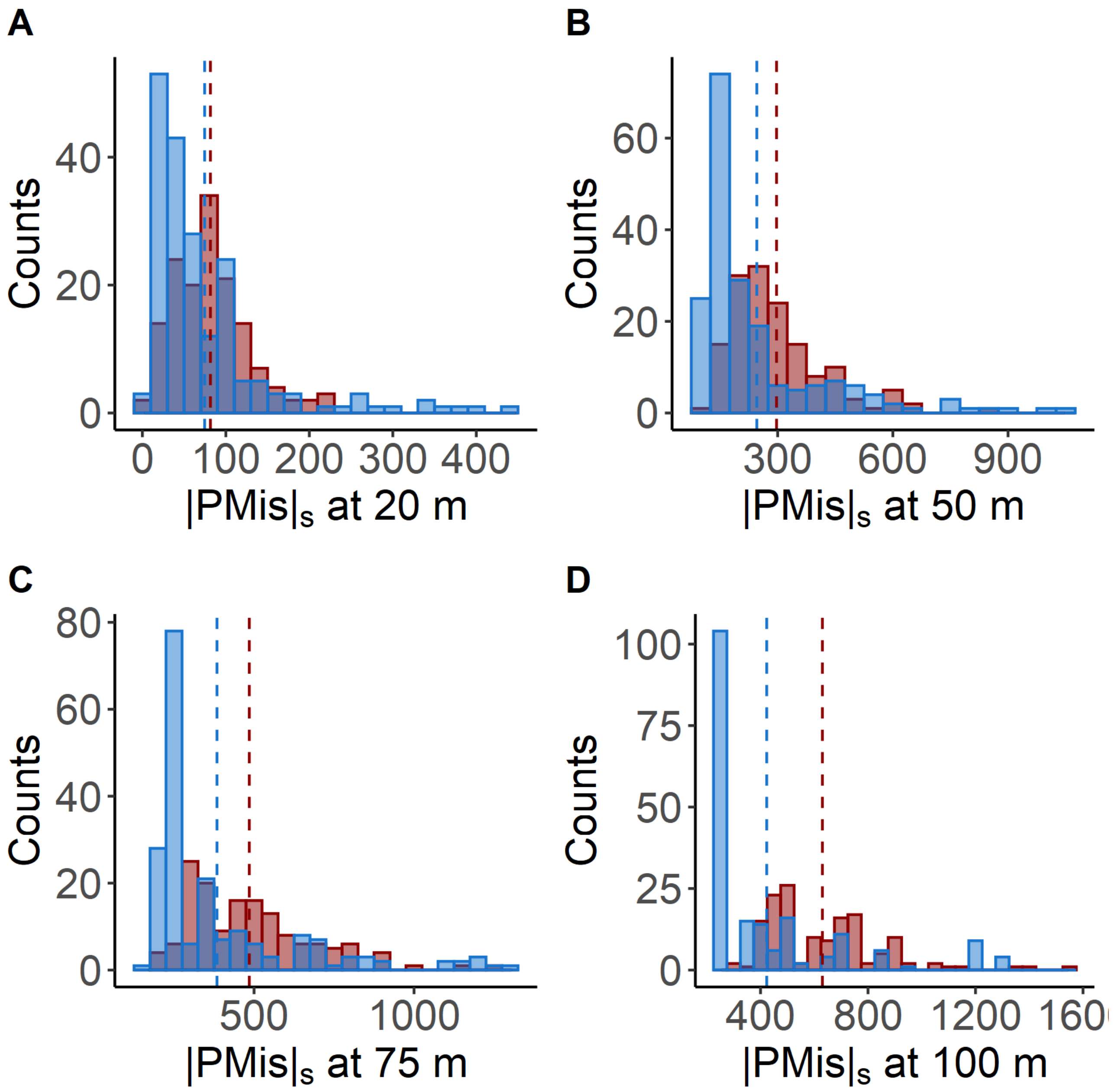

**Figure S11. Distribution of mean phenological mismatch (|Pmis|$_m$).**

|Pmis|$_m$ was computed as the mean absolute differences in TBB between a focal tree and its neighbors in various radius at plot N1-LOW (red bars) and N4-HIGH (blue bars). The dashed lines represent mean values in each plot, illustrating that |Pmis|$_m$ was always higher at low that at high altitude. See also the variance of |Pmis|$_m$ in Table S1.

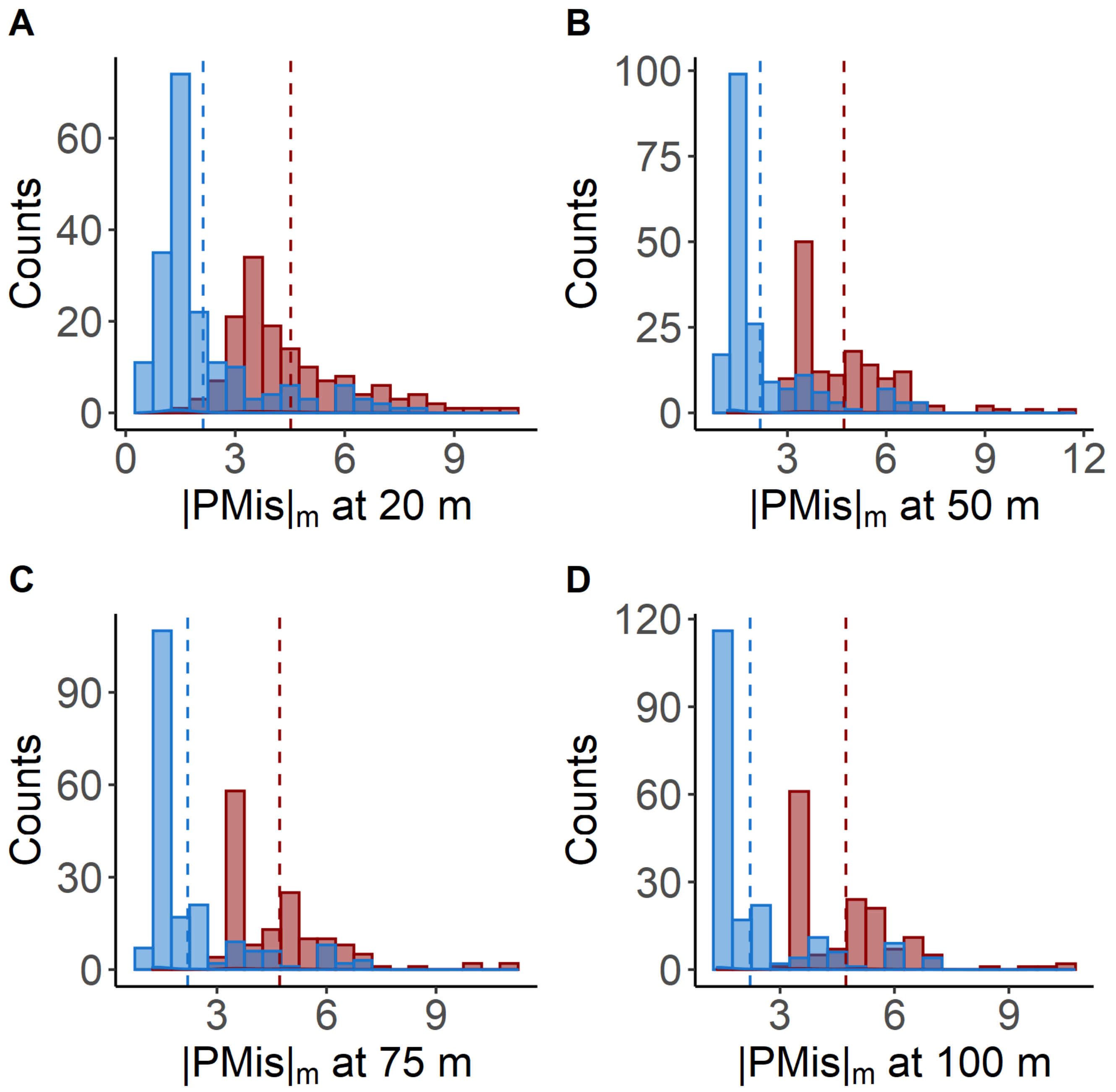

**Figure S12 Quality control of the fit of linear models for the estimation of sexual selection.**

*(A) Sexual selection model on female fecundity at plot N1-LOW (model M2.sex)*

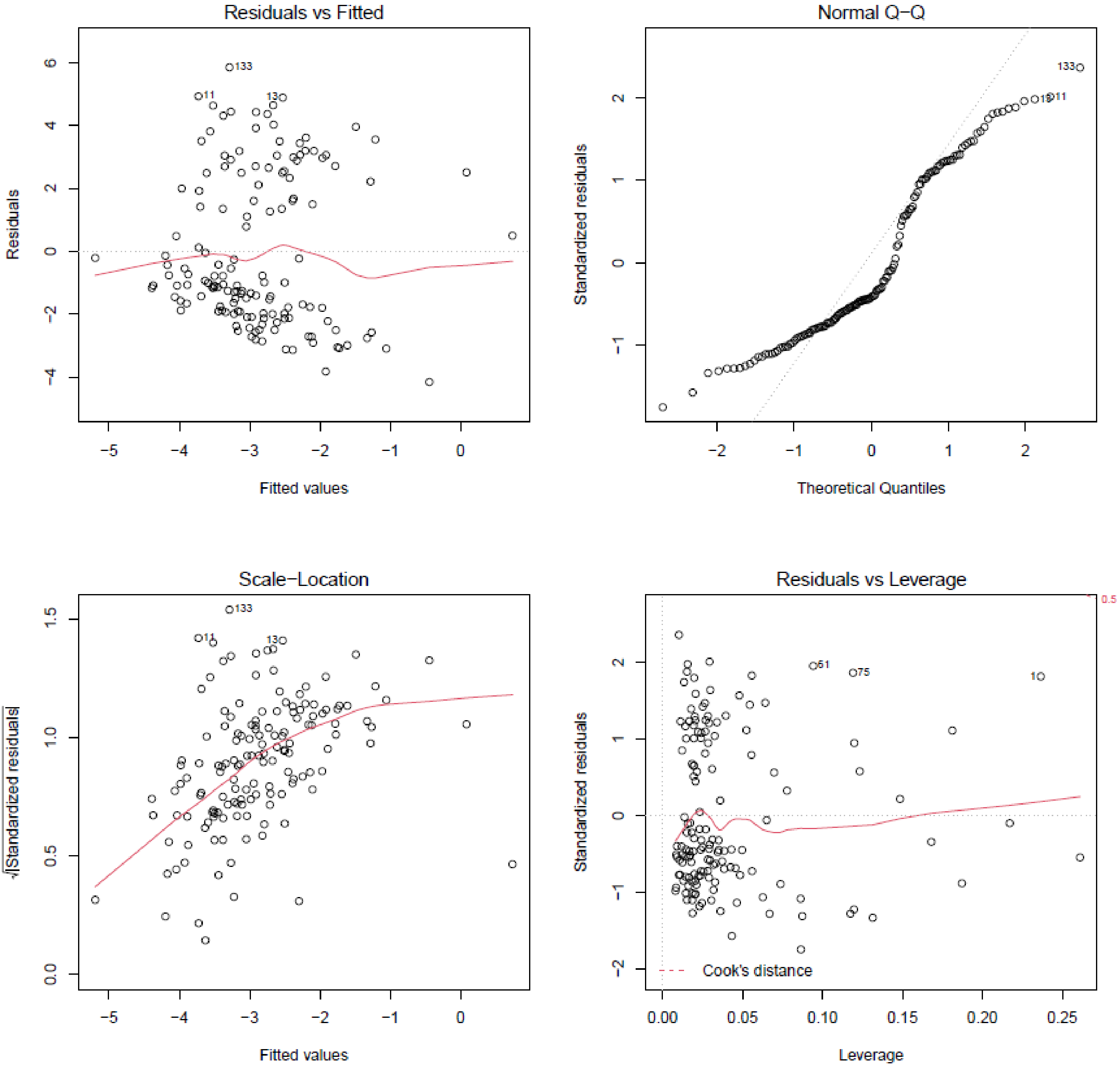

*(B) Sexual selection model on female fecundity at plot N4-HIGH (model M2.sex)*

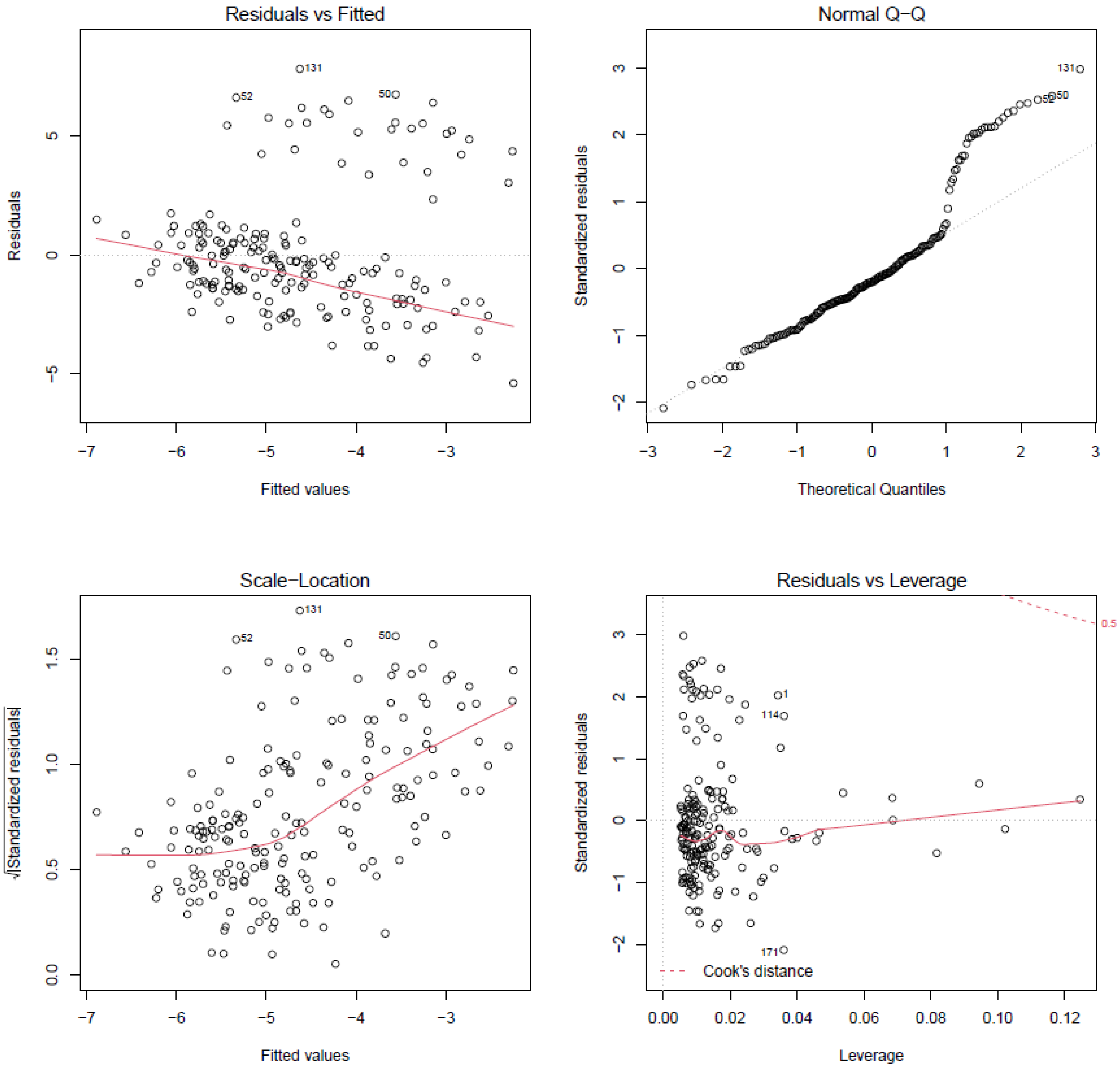

*(C) Sexual selection model on male fecundity at plot N1-LOW (model M2.sex)*

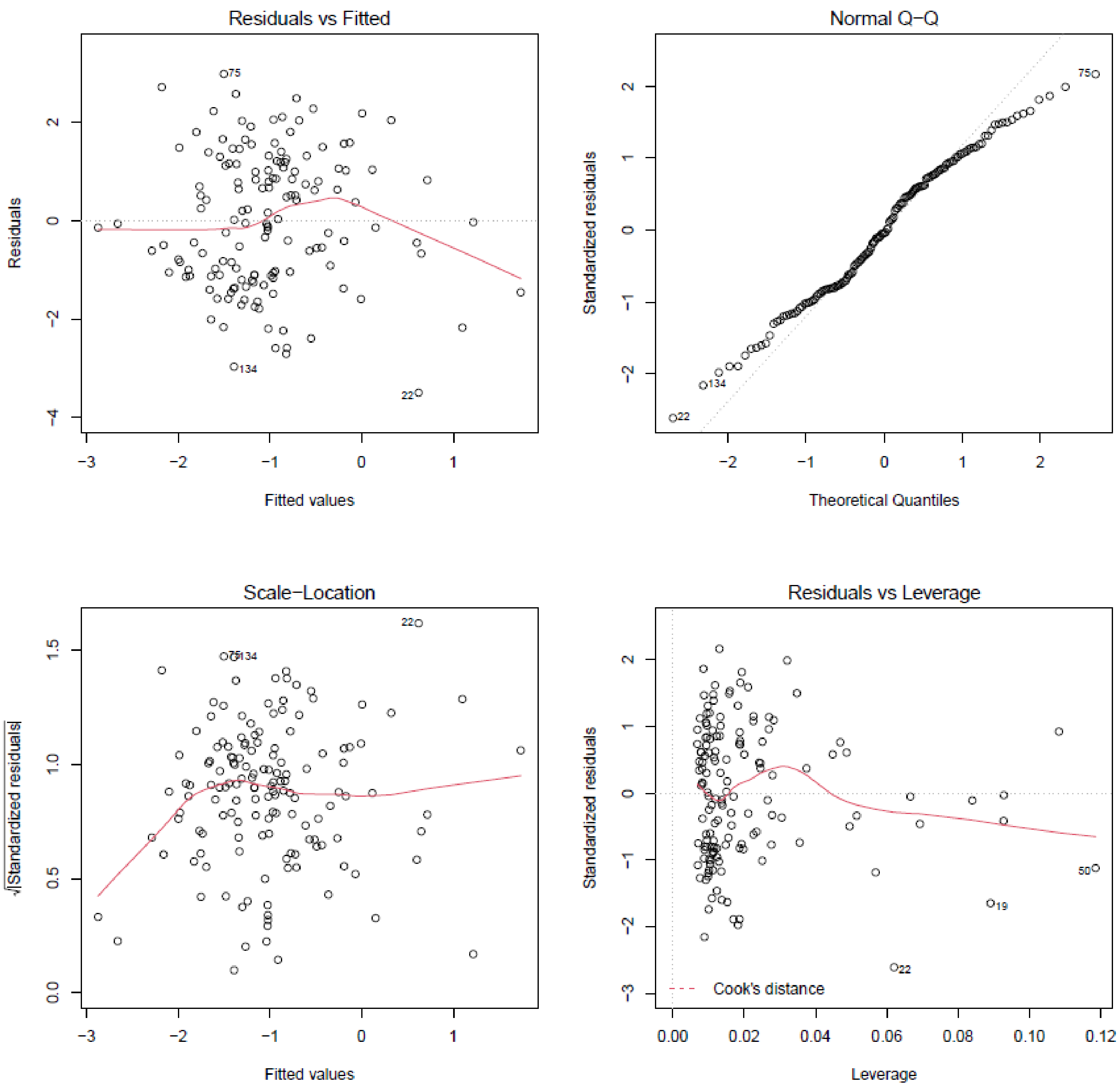

*(D) Sexual selection model on male fecundity at plot N4-HIGH (model M4.sex)*

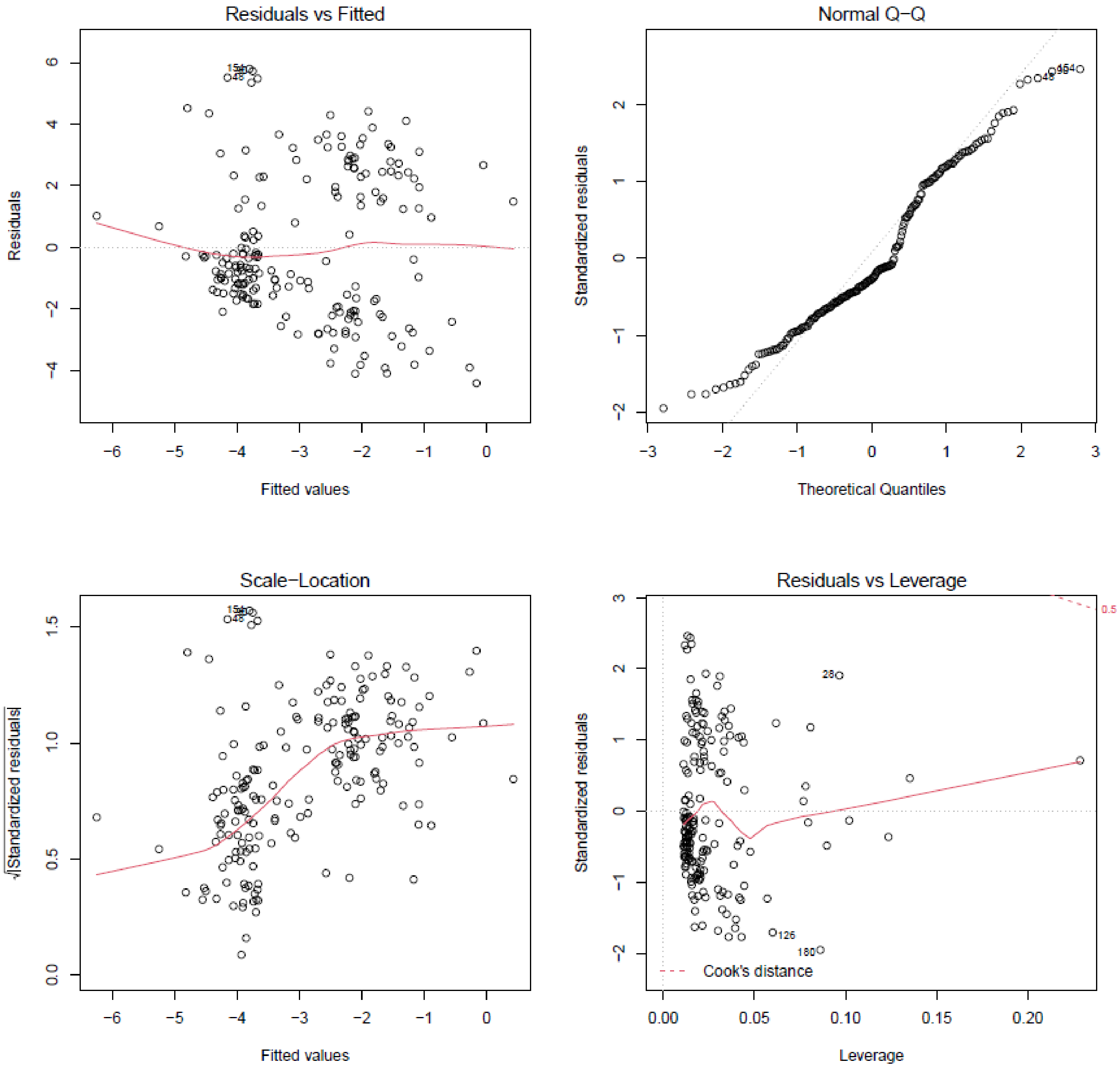

**Table S1: Variation of phenological mismatch estimators.**

Phenological mismatch was computed as the sum ($|PMis|_s$) or the mean ($|PMis|_m$) of differences in TBB between each focal tree and its neighbors in radius on 20, 50, 75 or 100 m.

| Radius | Plot | $\|PMis\|_s$ | | | $\|PMis\|_m$ | | |
|---|---|---|---|---|---|---|---|
| | | mean | sd | cv | mean | sd | cv |
| 20 | N1-LOW | 81.51 | 43.86 | 0.54 | 4.52 | 1.75 | 0.39 |
| | N4-HIGH | 74.56 | 77.95 | 1.05 | 2.12 | 1.50 | 0.71 |
| 50 | N1-LOW | 297.13 | 125.52 | 0.42 | 4.72 | 1.54 | 0.33 |
| | N4-HIGH | 245.69 | 177.40 | 0.72 | 2.17 | 1.44 | 0.66 |
| 75 | N1-LOW | 485.24 | 193.05 | 0.40 | 4.72 | 1.50 | 0.32 |
| | N4-HIGH | 384.17 | 244.88 | 0.64 | 2.20 | 1.42 | 0.64 |
| 100 | N1-LOW | 630.30 | 213.57 | 0.34 | 4.73 | 1.47 | 0.31 |
| | N4-HIGH | 423.05 | 270.20 | 0.64 | 2.22 | 1.41 | 0.64 |

**Table S2 Comparison of fecundity selection models on female and male fecundity at both plots.** For each model we display the number of parameters (K), the the Akaike information criterion corrected for small sample size (AICc), the difference in AICc between each model and the most parsimonious model (Δ AICc), and the log-likehood of the model (LL), all computed using the R package 'AICcmodavg'. The most parsimonious model is on the top line (grey). Codes and details in Online Appendix 1.

**A. Female fecundity, Plot N1-LOW.**

| Model name | K | AICc | Δ AICc | LL |
|---|---|---|---|---|
| **M7.fec (complete)** | 7 | 682.82 | 0.00 | -334.01 |
| **M5.fec (TBB+maxDBH+ConMar20+TBB:MAxDbh)** | 6 | 686.36 | 3.54 | -336.88 |
| **M2.fec (TBB+maxDbh)** | 4 | 686.57 | 3.75 | -339.15 |
| **M4.fec (TBB+maxDBH+ConMar20)** | 5 | 688.71 | 5.89 | -339.14 |
| **M6.fec (TBB+maxDBH+ConMar20+TBB:ConMar20)** | 6 | 690.11 | 7.29 | -338.75 |
| **M3.fec (TBB+ConMar20)** | 4 | 696.38 | 13.56 | -344.05 |
| **M1.fec (TBB)** | 3 | 698.05 | 15.23 | -345.94 |
| **Null** | 2 | 699.63 | 16.81 | -347.77 |

**B. Female fecundity, Plot N4-HIGH**

| Model | K | AICc | Δ AICc | LL |
|---|---|---|---|---|
| **M5.fec (TBB+maxDBH+ConMar20+TBB:MAxDbh)** | 6 | 917.07 | 0.00 | -452.31 |
| **M4.fec (TBB+maxDBH+ConMar20)** | 5 | 917.63 | 0.56 | -453.65 |
| **M7.fec (complete)** | 7 | 919.22 | 2.15 | -452.31 |
| **M6.fec (TBB+maxDBH+ConMar20+TBB:ConMar20)** | 6 | 919.35 | 2.29 | -453.45 |
| **M2.fec (TBB+maxDbh)** | 4 | 919.45 | 2.38 | -455.62 |
| **M3.fec (TBB+ConMar20)** | 4 | 927.13 | 10.06 | -459.46 |
| **M1.fec (TBB)** | 3 | 941.25 | 24.18 | -467.56 |
| **Null** | 2 | 942.83 | 25.76 | -469.38 |

**C. Male fecundity, Plot N1-LOW**

| Model | K | AICc | Δ AICc | LL |
|---|---|---|---|---|
| **M2.fec (TBB+sumDbh)** | 4 | 531.46 | 0.00 | -261.59 |
| **M4.fec (TBB+sumDbh+ConDens20)** | 5 | 532.05 | 0.59 | -260.81 |
| **M5.fec (TBB+sumDbh+ConDens20+TBB:sumDbh)** | 6 | 532.22 | 0.75 | -259.81 |
| **M7.fec (complete)** | 7 | 532.55 | 1.09 | -258.87 |
| **M6.fec (TBB+sumDbh+ConDens20+TBB:ConDens20)** | 6 | 533.28 | 1.81 | -260.34 |
| **Null** | 2 | 547.90 | 16.43 | -271.91 |
| **M1.fec (TBB)** | 3 | 549.88 | 18.41 | -271.85 |
| **M3.fec (TBB+ConDens20)** | 4 | 551.99 | 20.53 | -271.85 |

**D. Male fecundity, Plot N4-HIGH**

| Model | K | AICc | Δ AICc | LL |
|---|---|---|---|---|
| **M4.fec (TBB+sumDbh+Stature)** | 6 | 884.12 | 0.00 | -435.83 |
| **M5.fec (TBB+sumDbh+Stature+TBB:sumDbh)** | 7 | 886.25 | 2.13 | -435.82 |
| **M6.fec (TBB+sumDbh+Stature+TBB:Stature)** | 8 | 887.65 | 3.52 | -435.43 |
| **M3.fec (TBB+ConDens20)** | 5 | 888.03 | 3.91 | -438.85 |
| **M7.fec (complete)** | 9 | 889.70 | 5.57 | -435.35 |
| **M2.fec (TBB+sumDbh)** | 4 | 896.69 | 12.56 | -444.24 |
| **M1.fec (TBB)** | 3 | 911.60 | 27.48 | -452.74 |
| **Null** | 2 | 916.97 | 32.85 | -456.45 |

**Table S3 Comparison of sexual selection models on female and male fecundity at both plots.**
For each model we display the number of parameters (K), the the Akaike information criterion corrected for small sample size (AICc), the difference in AICc between each model and the most parsimonious model (Δ AICc), and the log-likehood of the model (LL), all computed using the R package 'AICcmodavg'. The most parsimonious model is on the top line (grey). Codes and details in Online Appendix 1.

**A. Female fecundity, Plot N1-LOW.**

| Model | K | AICc | Δ AICc | LL |
|---|---|---|---|---|
| **M2.sex (Pmis+maxDbh)** | 4 | 688.31 | 0.00 | -340.01 |
| **M4.sex (Pmis+maxDBH+ConMar20)** | 5 | 690.43 | 2.12 | -340.00 |
| **M5.sex (Pmis+maxDBH+ConMar20+Pmis:MAxDbh)** | 6 | 692.56 | 4.25 | -339.98 |
| **M6.sex (Pmis+maxDBH+ConMar20+Pmis:ConMar20)** | 6 | 692.57 | 4.26 | -339.99 |
| **M7.sex (complete)** | 7 | 693.56 | 5.25 | -339.38 |
| **Null** | 2 | 699.63 | 11.32 | -347.77 |
| **M3.sex (Pmis+ConMar20)** | 4 | 699.74 | 11.43 | -345.73 |
| **M1.sex (Pmis)** | 3 | 701.41 | 13.10 | -347.62 |

**B. Female fecundity, Plot N4-HIGH**

| Model | K | AICc | Δ AICc | LL |
|---|---|---|---|---|
| **M2.sex (Pmis+maxDbh)** | 4 | 921.00 | 0.00 | -456.39 |
| **M4.sex (Pmis+maxDBH+ConMar20)** | 5 | 921.31 | 0.32 | -455.49 |
| **M5.sex (Pmis+maxDBH+ConMar20+Pmis:MAxDbh)** | 6 | 922.66 | 1.66 | -455.10 |
| **M6.sex (Pmis+maxDBH+ConMar20+Pmis:ConMar20)** | 6 | 923.16 | 2.17 | -455.36 |
| **M7.sex (complete)** | 7 | 924.59 | 3.60 | -454.99 |
| **M3.sex (Pmis+ConMar20)** | 4 | 930.38 | 9.39 | -461.09 |
| **M1.sex (Pmis)** | 3 | 942.14 | 21.14 | -468.00 |
| **Null** | 2 | 942.83 | 21.83 | -469.38 |

**C. Male fecundity, Plot N1-LOW**

| Model | K | AICc | Δ AICc | LL |
|---|---|---|---|---|
| **M2.sex (Pmis+sumDbh)** | 4 | 517.48 | 0.00 | -254.60 |
| **M4.sex (Pmis+sumDbh+ConDens20)** | 5 | 517.54 | 0.05 | -253.56 |
| **M6.sex (Pmis+sumDbh+ConDens20+Pmis:ConDens20)** | 6 | 517.69 | 0.20 | -252.54 |
| **M5.sex (Pmis+sumDbh+ConDens20+Pmis:sumDbh)** | 6 | 519.22 | 1.73 | -253.31 |
| **M7.sex (complete)** | 7 | 520.32 | 2.84 | -252.76 |
| **M3.sex (Pmis+ConDens20)** | 4 | 537.66 | 20.17 | -264.69 |
| **M1.sex (Pmis)** | 3 | 541.30 | 23.81 | -267.57 |
| **Null** | 2 | 547.90 | 30.41 | -271.91 |

**D. Male fecundity, Plot N4-HIGH**

| Model | K | AICc | Δ AICc | LL |
|---|---|---|---|---|
| **M4.comp (Pmis+TBB+sumDbh+Stature)** | 7 | 881.06 | -1.99 | -433.23 |
| **M4.sex (Pmis+sumDbh+Stature)** | 6 | 883.05 | 0.00 | -435.30 |
| **M5.sex (Pmis+sumDbh+Stature+Pmis:sumDbh)** | 7 | 884.40 | 1.35 | -434.89 |
| **M6.sex (Pmis+sumDbh+Stature+Pmis:Stature)** | 8 | 885.01 | 1.96 | -434.11 |
| **M3.sex (Pmis+Stature)** | 5 | 886.04 | 2.98 | -437.86 |
| **M7.sex (complete)** | 10 | 887.01 | 3.95 | -432.90 |
| **M2.sex (Pmis+sumDbh)** | 4 | 899.26 | 16.21 | -445.52 |
| **M1.sex (Pmis)** | 3 | 914.45 | 31.40 | -454.16 |
| **Null** | 2 | 916.97 | 33.92 | -456.45 |

**Table S4 Comparison of best models for fecundity and sexual selection with compound models.**
For each plot and each sex, we tested compound best models, including both TBB and Pmis as factors , and compared them with the best models for fecundity and sexual selection identified in Table S2 and TableS3.For each model we display the number of parameters (K), the Akaike information criterion corrected for small sample size (AICc), the difference in AICc between each model and the most parsimonious model (Δ AICc), and the log-likehood of the model (LL), all computed using the R package 'AICcmodavg'. The most parsimonious model is on the top line (grey). Codes and details in Online Appendix 1.

| Plot | Sex | Model name (factors included) | K | AICc | Delta_AICc | LL |
|---|---|---|---|---|---|---|
| N1-LOW | Female | M7.fec (complete, TBB) | 7 | 682.82 | 0.00 | -334.01 |
| | | M7.comp (complete, Pmis+TBB) | 8 | 684.84 | 2.02 | -333.90 |
| | | M7.sex (complete, Pmis) | 7 | 693.56 | 10.74 | -339.38 |
| N4-HIGH | Female | M5.fec (TBB+maxDBH+ConMar20+…) | 6 | 917.07 | 0.00 | -452.31 |
| | | M5.comp (idem, with Pmis+TBB) | 7 | 919.20 | 2.13 | -452.30 |
| | | M5.sex (idem, Pmis) | 6 | 922.66 | 5.59 | -455.10 |
| N1-LOW | Male | M2.sex (Pmis+sumDbh) | 4 | 517.48 | 0.00 | -254.60 |
| | | M2.comp (TBB+Pmis+sumDbh) | 5 | 519.47 | 1.99 | -254.52 |
| | | M2.fec (TBB+sumDbh) | 4 | 531.46 | 13.98 | -261.59 |
| N4-HIGH | Male | M4.comp (TBB+Pmis+sumDbh+Stature) | 7 | 881.06 | 0.00 | -433.23 |
| | | M4.sex (Pmis+sumDbh+Stature) | 6 | 883.05 | 1.99 | -435.30 |
| | | M4.fec (TBB+sumDbh+Stature) | 6 | 884.12 | 3.06 | -435.83 |

**Table S5 Standardized selection gradients (β') on phenological traits with their standard deviation (σ) for female and male fecundity at both plots.**

Standardized selection gradients were estimated using the best models selected for fecundity and sexual selection (see Table S2 and S3 above). These models already included standardized traits, but were fitted on log-transformed fecundity estimates. We refitted them on non-transformed fecundity estimates to obtain the selection gradients (β') and their standard errors (s.e.) displayed in this table.

| Sex | Plot | Trait | Model | σ | β' | s.e. |
|---|---|---|---|---|---|---|
| **Female** | N1-LOW | TBB | M7.fec | 4.160 | -0.24 | 0.19 |
| | N4-HIGH | TBB | M5.fec | 2.286 | -0.43 | 0.25 |
| **Male** | N1-LOW | \|PM\| | M2.sex | 43.865 | -0.40 | 0.12 |
| | N4-HIGH | TBB | M4.comp | 2.286 | -0.30 | 0.16 |
| | N4-HIGH | \|PM\| | M4.comp | 77.95 | -0.16 | 0.16 |

**Table S6 Comparison of viability selection models on seedlings growth in diameter (DGrowth) and height (HGrowth).**

For each response variable (DGrowth and HGrowth), we fitted several models: M1viabSel and M2viabSel are detailed in the main text. In the model “M1viabSel – Block” the random effect Block was removed (to test for the significance of Block effect). In the model “M1viabSel – Family” the random effect Family was removed (to test for the significance of Family effect). For each model we display the number of parameters (K), the Akaike information criterion (AIC), the log-likehood of the model (LL). Codes and details in Online Appendix 2.

| | K | AIC | LL | Deviance | $Chi^2$ | Df | Pr(>$Chi^2$) |
|---|---|---|---|---|---|---|---|
| DGrowth | | | | | | | |
| M1viabSel - Block | 7 | 7777.9 | -3881.9 | 7763.9 | | | |
| M1viabSel - Family | 7 | 7853.9 | -3920 | 7839.9 | 0 | 0 | |
| M1viabSel | 8 | 7753 | -3868.5 | 7737 | 102.93 | 1 | < 2.20 10-16 |
| M2viabSel | 12 | 7735.8 | -3855.9 | 7711.8 | 25.181 | 4 | 4.63E-05 |
| HGrowth | | | | | | | |
| M1viabSel - Block | 7 | 35588 | -17787 | 35574 | | | |
| M1viabSel - Family | 7 | 35734 | -17860 | 35720 | 0 | 0 | |
| M1viabSel | 8 | 35544 | -17764 | 35528 | 191.36 | 1 | < 2.20 10-16 |
| M2viabSel | 12 | 35478 | -17727 | 35454 | 74.693 | 4 | 2.31E-15 |

**Table S7: Viability selection on seedling growth in diameter (Dgrowth) and height (Dgrowth).** Selection on phenology was assessed through the effect of TBB on seedling growth (A-Dgrowth and B-Hgrowth), accounting for the effects of plot (N1-LOW or N4-HIGH), initial size (initD or initH), Treatment (TTmt, i.e., Stressed- Str- versus Watered- Wat) and common garden design (with Block and Family included as random effects). Significance of the effect (*p*-values) was assessed based on the t-value and dof.

**A Diameter growth**

| Term | K | SoSq | F-value | p-value | Effect | se | t-value | p-value |
|---|---|---|---|---|---|---|---|---|
| Plot | 1 | 2.78 | 4.03 | 0.045 | -0.34 | 0.386 | -0.879 | |
| TTmt | 1 | 10.20 | 14.82 | <0.001 | -1.60 | 0.381 | -4.190 | |
| initD | 1 | 222.18 | 322.61 | <0.001 | 0.29 | 0.016 | 18.381 | |
| Plot:TTmt | 1 | 0.32 | 0.47 | 0.494 | -0.24 | 0.377 | -0.645 | |
| Plot:TTmt:TBB | 4 | 29.33 | 10.65 | <0.001 | N1-LOW, Wat: 0.00 | 0.008 | 0.523 | 0.601 |
| | | | | | N4-HIGH, Wat: -0.01 | 0.008 | -0.778 | 0.437 |
| | | | | | N1-LOW, Str: -0.03 | 0.008 | -4.417 | <0.001 |
| | | | | | N4-HIGH, Str: -0.03 | 0.007 | -4.750 | <0.001 |

**B Height growth**

| Term | K | SoSq | F-value | p-value | Effect | se | t-value | p-value |
|---|---|---|---|---|---|---|---|---|
| Plot | 1 | 67518 | 11.86 | <0.001 | 7.80 | -35.207 | 2.210 | |
| TTmt | 1 | 229800 | 40.37 | <0.001 | -94.09 | 34.636 | -2.717 | |
| initH | 1 | 18644 | 3.28 | 0.070 | 0.05 | 0.019 | 2.344 | |
| Plot:TTmt | 1 | 11087 | 1.95 | 0.163 | -5.16 | 34.276 | -0.151 | |
| Plot:TTmt:TBB11 | 4 | 116660 | 5.12 | 0.000 | N1-LOW, Wat: 0.99 | 0.712 | 1.394 | 0.163 |
| | | | | | N4-HIGH, Wat: -0.43 | 0.722 | -0.591 | 0.555 |
| | | | | | N1-LOW, Str: -1.26 | 0.691 | -1.828 | 0.068 |
| | | | | | N4-HIGH, Str: -2.59 | 0.673 | -3.852 | <0.001 |
